# Macromolecular interactions and geometrical confinement determine the 3D diffusion of ribosome-sized particles in live *Escherichia coli* cells

**DOI:** 10.1101/2024.03.27.587083

**Authors:** Diana Valverde-Mendez, Alp M. Sunol, Benjamin P. Bratton, Morgan Delarue, Jennifer L. Hofmann, Joseph P. Sheehan, Zemer Gitai, Liam J. Holt, Joshua W. Shaevitz, Roseanna N. Zia

## Abstract

The crowded bacterial cytoplasm is comprised of biomolecules that span several orders of magnitude in size and electrical charge. This complexity has been proposed as the source of the rich spatial organization and apparent anomalous diffusion of intracellular components, although this has not been tested directly. Here, we use biplane microscopy to track the 3D motion of self-assembled bacterial Genetically Encoded Multimeric nanoparticles (bGEMs) with tunable size (20 to 50 nm) and charge (−2160 to +1800 e) in live *Escherichia coli* cells. To probe intermolecular details at spatial and temporal resolutions beyond experimental limits, we also developed a colloidal whole-cell model that explicitly represents the size and charge of cytoplasmic macromolecules and the porous structure of the bacterial nucleoid. Combining these techniques, we show that bGEMs spatially segregate by size, with small 20-nm particles enriched inside the nucleoid, and larger and/or positively charged particles excluded from this region. Localization is driven by entropic and electrostatic forces arising from cytoplasmic polydispersity, nucleoid structure, geometrical confinement, and interactions with other biomolecules including ribosomes and DNA. We observe that at the timescales of traditional single molecule tracking experiments, motion appears sub-diffusive for all particle sizes and charges. However, using computer simulations with higher temporal resolution, we find that the apparent anomalous exponents are governed by the region of the cell in which bGEMs are located. Molecular motion does not display anomalous diffusion on short time scales and the apparent sub-diffusion arises from geometrical confinement within the nucleoid and by the cell boundary.

## INTRODUCTION

In the absence of membrane bound-compartments and transport motor proteins [1, 2], bacterial cells rely almost exclusively on thermal, electrostatic, and hydrodynamic interactions to achieve spatial organization that is important for biological function [3, 4, 5, 6, 7, 8]. These interactions take place within the extremely crowded and polydisperse bacterial cytoplasm [9, 10, 11], whose components span multiple orders of magnitude in both size and charge. Charge and size are typically linked, with ions of size *∼* 0.1 nm being weakly positive (*∼* 1*e*), proteins of size *∼* 1 nm roughly neutral to weakly negative (*∼* 1e - 10 e), and large macromolecular complexes such as ribosomes, polysomes and nucleic acids having large negative charges (*∼* 1, 000 *−* 10, 000*e*) [12]. The bacterial chromosome is condensed near the center of the cell in a highly organized structure called the nucleoid [12, 13]. In *E. coli* the nucleoid occupies 35-65% of the cellular volume [14, 15, 16, 17], with a mesh size of *≈* 50 nm [18]. Thus, large macromolecular complexes, such as active ribosomes and polysomes, tend to be excluded from this region [19, 20, 13, 21, 22, 23], while ribosomal sub-units and small proteins are free to penetrate into and diffuse within [13, 21, 24, 25] the membrane-less nucleoid. Electrostatic charge plays an important role in spatial segregation of the nucleoid [12] and intermolecular interactions outside the nucleoid [26, 27]. However, its effect on intracellular particle localization is unknown.

Beyond this static picture of intracellular organization, the bacterial cytoplasm is a highly dynamic environment. Intracellular search and transport processes crucial for cell growth and survival are primarily driven by thermal diffusion [28, 29, 30], but the size- and charge-dependent dynamics of molecules in the cytoplasm are not fully understood. Previous single-particle tracking (SPT) experiments of molecules have reported sub-diffusion in living bacterial cells [31, 22, 16, 32, 33], which might play an important role in bacterial regulation by keeping molecules close to their target sites and reducing search times [34]. For small particles, such as proteins (*∼* 1 nm), any apparent sub-diffusivity can be attributed to the effects of confinement within the small cellular volume (*∼* 0.1-1 *µ*m^3^) [35, 36]. For larger particles (*∼* 10 nm - *∼* 100 nm), both diffusive [18] and sub-diffusive behavior [31, 22, 16, 32, 33] have been reported. This slow down in particle dynamics has been proposed to emerge from cytoplasmic complexity instead of geometrical effects [34, 31, 33]. The proposed mechanisms include: caging due to extreme crowding [37, 34, 38], binding and unbinding events [34], spatial and dynamic heterogeneities in the cytoplasm [33], and non-ergodicities of a “glassy bacterial cytoplasm” [31]. However, there is no clear consensus on the degree of anomalous diffusion or the primary mechanism driving this behavior. Furthermore, little is known about the effects of particle charge on dynamics. It has been shown that positive proteins diffuse considerably slower than their negative counterparts [39, 19], but we lack an understanding of how charge affects the motion of larger macromolecules.

Previous SPT experiments have been unable to directly address these questions due to limitations in dimensionality and spatio-temporal resolution. Studies reporting sub-diffusion have relied on two-dimensional (2D) SPT measurements. However, interpreting the actual 3D motion from 2D projections is challenging [40]. Typical observed trajectories are short such that statistical analyses of the displacement only cover approximately one order of magnitude in time scale. Finally, some strategies for generating macromolecular sized particles for tracking result in polydisperse assemblies[34, 31], precluding accurate control of the characteristics such as size and charge.

Physics-based modeling of the bacterial cytoplasm serves as a promising avenue to complement SPT experiments by interrogating microscopic forces and fluid dynamics at shorter length and timescales than can be achieved experimentally [11, 41]. Current models of diffusion in the bacterial cytoplasm focused on the nucleoid-free region [9, 42, 41, 26, 27], neglecting both confinement [43, 44, 45] and the nucleoid, which are essential to study dynamics and organization on cellular length scales. Dynamic simulations of the nucleoid itself [46, 47, 48, 49, 50] have not represented the size-polydispersity of the rest of the crowded cytoplasm, and therefore are not easily adaptable to study more general features of heterogeneous macromolecular composition and cellular-scale dynamics in *E. coli*.

In this study, we aim to understand the role of particle charge and size on *in vivo* dynamics, considering different attributes of the heterogeneous intracellular environment across multiple timescales, including confinement, size polydispersity, intermolecular interactions, and the presence of a porous nucleoid. To do so, we combine high spatial and temporal resolution 3D-SPT of bacterial Genetically Encoded Multimeric Nanoparticles (bGEMs) of tunable fixed diameter (20, 40, and 50 nm) and charge (−2160, −840, +1800*e*) with colloidal whole-cell dynamical modeling as well as longer-time probabilistic simulations of an excluded volume nucleoid to interrogate how macromolecular and cell-scale characteristics affect emergent dynamic behaviors in *E. coli*. Our results show that size and charge primarily determine where bGEMs localize inside the cell. This localization emerges from entropic and electrostatic interactions of the particle with the nucleoid pore structure and the rest of the cytoplasmic milieu. Importantly, we find that confinement effects can primarily explain the anomalous diffusion measured in previous SPT studies. Combining 3D-SPT experiments and coarse-grained whole-cell modeling enables us to explore dynamics below the temporal resolution of 3D-SPT and suggests physical mechanisms underlying the experimentally observed phenomena.

## RESULTS

We use biplane microscopy to track the 3D motion of self-assembled nanoparticles, each of a fixed size and charge in live *E.coli* cells. We adapted these bacterial nanoparticles, termed bGEMs, from eukaryotic Genetically Encoded Multimeric nanoparticles (GEMs) [55]. Biplane microscopy enables us to image simultaneously two focal planes, resulting in three-dimensional information of the particle’s position with sub-pixel resolution (see Table S1 for localization errors) and the ability to track single particles for hundreds of frames (*∼* 1 s) at a rate of 30 ms per frame (SI Movie 1). Until now, this time resolution has been absent in the bacterial literature. A typical experimental data set is exemplified in Figure 1(a-c) and SI Movie 2. Three representative time frames from a bacterial cell are shown, with the DNA labeled in magenta, and the cellular wall labeled in blue. Typically, a single bGEM (green) is self-assembled per cell. We use custom software (See S1 in SI) to reconstruct each particle’s trajectory in 3D (Fig. 1(b), SI Movie 3). When measuring and describing 3D trajectories, we adopt a right-handed coordinate system such that the long axis of the cell is aligned along *x* and the focal axis of the microscope is aligned along *z*. Figure 1(c) shows the *yz* cross-section of the trajectory in (b). This cross-section clearly exemplifies nucleoid exclusion, with the particle avoiding almost entirely the interior of the cell. Inferring this type of information from a 2D-SPT projection would be challenging as it would require extensive probabilistic modeling to reconstruct a likely 3D trajectory. We chose to focus our analysis in stationary phase cells to avoid growth effects and to preserve the shape of the nucleoid. However, we observe excellent agreement between exponential and stationary phase localization and dynamics (Fig. S1).

**Figure 1.**
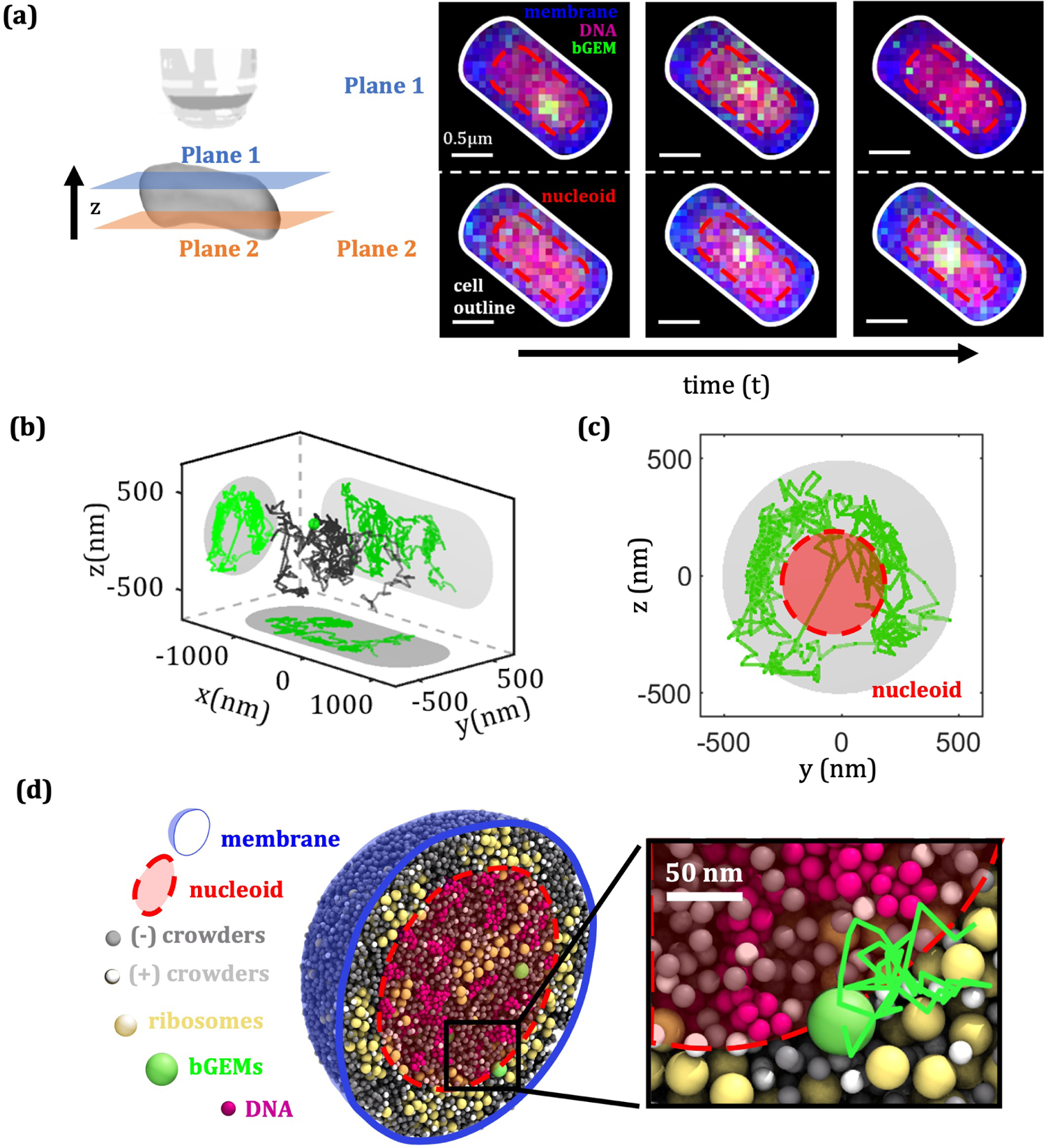
3D-SPT experiments and whole-cell colloidal simulations probe macromolecular dynamics in E. coli. (a) Sample bacterial cell imaged through biplane microscopy. In biplane microscopy two focal planes are imaged simultaneously (schematic, left). Three snapshots are shown (images, right) from left to right in Plane 1 (top row) and Plane 2 (bottom row). The bacterial membrane is labeled in dark blue and the DNA is stained in magenta. Cell and nucleoid envelopes are outlined in white and red, respectively. The bGEM nanoparticle (green) is initially in focus in Plane 1 (top row, left); as it moves through the cell, it comes into focus on Plane 2 (bottom row, right). (b) Sample 3D trajectory for a 50nm particle. (c) yz projection of the trajectory shows the particle is primarily excluded from the cell’s nucleoid. (d) Whole-cell colloidal model of E. coli, cross-sectional view, dynamic simulation snapshot. The nucleoid (magenta, outlined in red) is interpenetrated and surrounded by a polydisperse cytoplasm, confined by a cellular membrane (blue). The cytoplasm consists of negatively (grey) and positively (white) charged native crowders (which represent proteins, ternary complexes, transcription factors, ribosomal subunits, etc), negatively charged ribosomes and polysomes (yellow), and bGEMs (green) at physiological abundances and densities [51, 52, 53, 41, 54]. The dynamical motion of individual bGEMs (green sphere, zoomed-in view, right) is tracked in simulation and compared to the 3D particle traces from experiments. Each particle moves via Brownian motion and interacts directly and physically with other macromolecules, the nucleoid, and the cell membrane.

We also developed a colloidal physics-based whole-cell model to investigate intermolecular dynamics at angstrom spatial and picosecond temporal resolutions, which are inaccessible to experiments (Fig. 1(d), SI Movie 4). Our model explicitly represent the size and charge polydispersity of macromolecules in the cytoplasm at physiological abundances derived from the literature and the porous structure of the bacterial nucleoid within a confining cell membrane (see Section and S4). This synergistic experimental and computational approach enables us to explore the impact of size and charge on diffusion inside *E. coli* from molecular to whole-cell length and time scales.

### Size-based particle segregation in *E. coli*

We first asked how macromolecular size affects sub-cellular particle localization inside the bacterial cell using three bGEM diameters: 20, 40, and 50 nm. In both experiment and simulation, we observe size-dependent localization of macromolecules in the bacterial cell. Figure 2 (a) shows 2D histograms of the position of detected particles in the *xy* and *yz* projection planes for each particle size. An outline of the average cell and nucleoid size are shown overlaid. The contrast of the images has been enhanced linearly to improve visibility. Raw images with a grayscale denoting number of localizations is shown in fig. S2 for all 3 projection planes (*xy*,*yz*,*xz*). The 20-nm bGEMs are enriched inside the interior nucleoid region, while the larger particles are preferentially localized in the cellular periphery with 40-nm particles exploring a larger fraction of the nucleoid than their 50-nm counterparts. These trends are evident in the radial distribution of the particle positions (Fig. 2 (b)), where the probability distributions shift towards the cellular periphery as particle size increases. The radial distributions are calculated exclusively for the cylindrical portion of the spherocylindrical cells because including the poles would bias the distributions towards the center of the cell. Therefore, since radial distributions do not capture enrichment at the poles, we also calculated the pole occupation (percentage of localizations compared to all localizations). For all particle sizes pole occupation is small, less than 15% (See Fig. S3). As expected, this number is smallest for the 20 nm particles, *≈* 5%, but unintuitively the pole occupation is highest for the mid-sized 40 nm particles (*≈* 15%) and not the 50 nm particles *≈* 10%. Poles account for close to 15% of the cellular volume, therefore this suggests that the 50 nm particles tend to localize at the periphery of the cylindrical segment of the cell and are excluded from the poles. Using custom code, we segmented the bacterial nucleoid for each cell using the DNA label shown in 1(a) and then classified the trajectory points as located inside or outside the nucleoid region, as shown in Fig. S10 (see S3 in SI for details). We calculated the nucleoid occupation time from this data, defined as the ratio of time spent inside the nucleoid over the total number of time points measured, and find that it decreases considerably with particle size (Fig. 2 (c)). We observe that 20-nm particles spend the majority of time inside the nucleoid (*≈* 60%) while the opposite is true for larger 40 and 50 nm particles (¡40%). Colloidal simulations replicate the experimental trends with excellent agreement as shown in 2(c).

**Figure 2.**
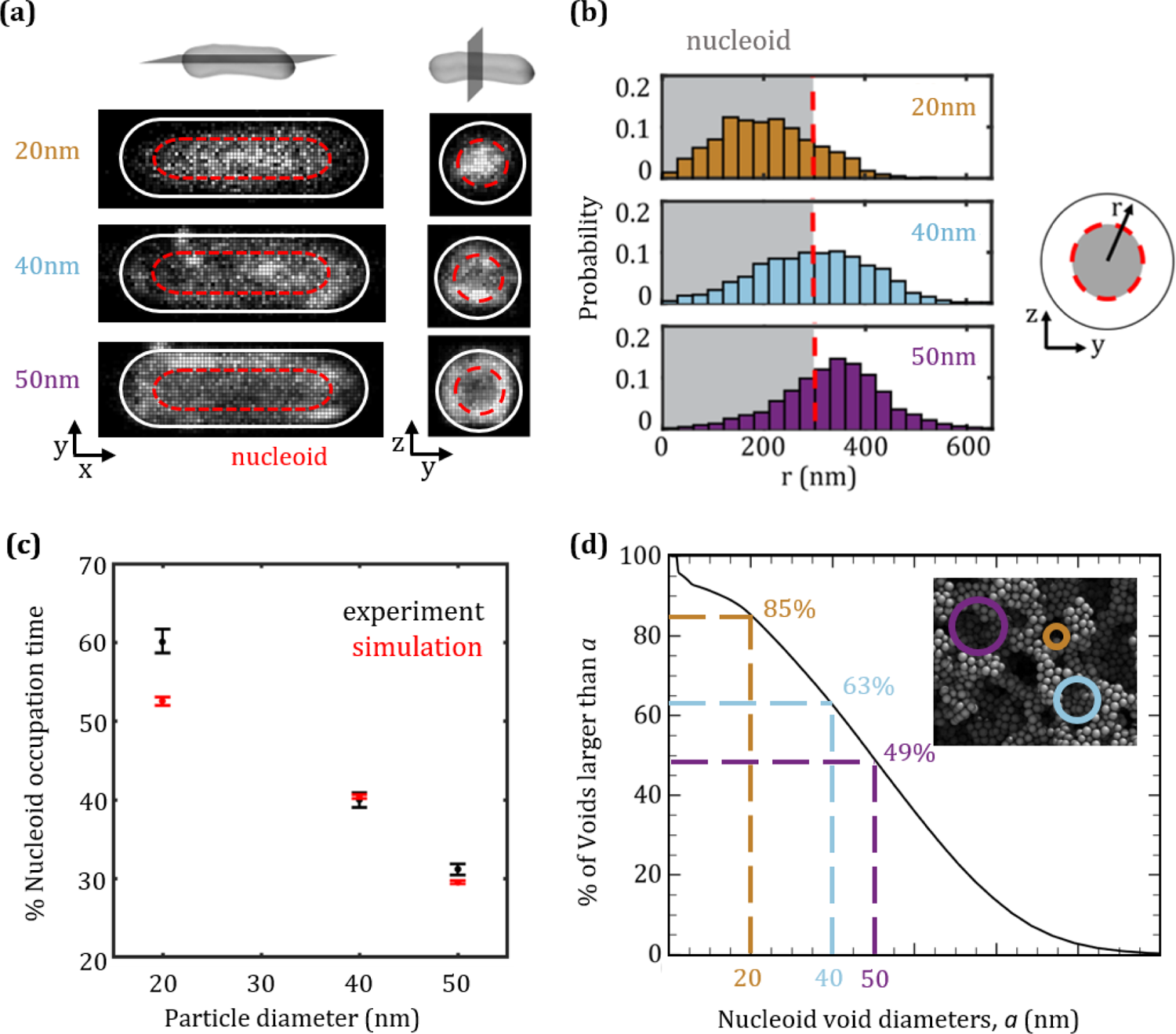
Macromolecular localization in E. coli depends on size. (a) *xy* and *yz* 2D histograms of20nm (3750 localizations), 40nm (9500 localizations), and 50nm (24000 localizations) tracers in top, middle, and bottom images, respectively. Cell sizes are normalized. Brighter intensity shows more localization. Contrast adjusted linearly to improve visibility. Average nucleoid region (red dashed line) and cell periphery (solid white line) are highlighted. (b) Radial density distributions (experiments) shift toward the cell periphery as tracer size increases. (c) Nucleoid occupation time (defined as total number of timepoints in the nucleoid divided by length of the trajectory) decreases with tracer size. The agreement between experiment (black) and simulations (red) is excellent. (d) Percent of voids in the nucleoid large enough to permit passage of a tracer of size *a*, for a range of tracer sizes shown on the horizontal axis. Inset: simulation snapshot of the interconnected nucleoid network with pores that permit penetration of a bGEM of size 20 nm (orange), 40 nm (light blue), and 50 nm (purple) highlighted. All bGEM sizes studied here can fit in some pores of the nucleoid, but smaller bGEMs (colored circles, inset) penetrate more of the void distribution.

We hypothesized that physical features of the nucleoid underlie the observed size-selective particle localization. To probe this hypothesis, we calculated the distribution of void (hole) sizes within the self-assembled nucleoid used in the whole-cell colloidal simulations. The microstructure of the porous network is heterogeneous: with an abundance of void sizes ranging from small *∼* 1 nm to large *∼* 100 nm (see Fig. S12). Using this distribution and void connectivity estimates for the simulated nucleoid, we calculated that particles with a diameter less than 80 nm would be able to diffuse into and pass through the entirety of the nucleoid. According to this theoretical prediction, all bGEM sizes should be able to fit within the pores of the nucleoid. However, the cumulative void size distributions shows that the percentage of accessible voids drops rapidly with particle size (Fig. 2 (d)). In particular, we found that 85%, 63%, and 49% of the voids are bigger than 20 nm, 40 nm, and 50 nm, respectively (dashed colored lines). The ratio of these percentages is close to the ratio of nucleoid occupation for the different sized bGEMs, suggesting that the underlying structure of the nucleoid’s empty space is responsible for the size-based spatial segregation we observe.

The lack of available space for larger molecules in the porous nucleoid structure results in their localization to the intracellular space outside the nucleoid. It is known that large macromolecules in *E. coli* such as active ribosomes and polysomes are excluded from the nucleoid [18]. This result is recapitulated in our model, where the equilibrium ribosome and polysome volume fraction outside the nucleoid is over 2.5 times higher than ribosomes inside the nucleoid. More specifically, the equilibrium ratio of single ribosome volume fraction outside to inside the nucleoid is approximately 0.95, whereas polysomal ribosomes (part of chains of six ribosomes) are almost completely excluded. These results are consistent with our experimentally-observed nucleoid occupation of bGEMs.

The exclusion of larger molecules from the nucleoid drives a small increase of packing fraction in the cytoplasm compared to that in the nucleoid (about 3%). The corresponding higher osmotic pressure in the cytoplasm tends to push smaller particles towards the center of the cell. This size segregation of particles in the cytoplasm is an example of entropic de-mixing, previously identified by Gonzalez et al. in confined colloidal suspensions [44]. We explored these size exclusion and demixing effects in a prior study of a simpler model cell, exploring strictly the role played by size variation and excluded volume effects in localization in and around a model nucleoid [56]. In that work, we showed that in the simpler monodisperse system, particles are preferentially excluded from the nucleoid, regardless of DNA compactness and density. But we found that when two or more particle sizes are present, the nucleoid becomes enriched in smaller particles whilst the surrounding cytoplasm becomes enriched in larger biomolecules, and the concentration of small molecules in the nucleoid is sometimes even higher than the nucleoid-free region. These simplified systems suggest that the localization trends in Figure 2 are a direct result of the cell’s broad polydispersity. The nucleoid’s void size distribution acts as an entropic size filter, enhancing the localization of small particles inside the DNA rich area and excluding larger particles to the cellular periphery. Our results are consistent with a nucleoid mesh size of *≈* 50 nm, in agreement with previous reports [18].

### Charge-based particle segregation in *E. coli*

In addition to the entropic interactions explored above, other forces present inside the cell may also affect particle localization. In particular, intermolecular, solvent-mediated electrostatic interactions are likely to alter particle position. In the size-effect studies, we used the msfGFP construct which carries a charge of *−*7*e* per protein. Using this fluorophore, 40 nm particles have a total charge of *−*840*e* and explore most of the cellular volume in both experiments and simulations. This diameter is thus ideal to observe charge-based localization effects. We expressed 40-nm bGEMs with GFP variants that produce total charges of *−*2, 160*e* and +1, 800*e*, allowing us to explore the effect of large negative and positive charges. We observe significant differences in particle localization between the charge-variant bGEMs, although this effect is smaller than for different sizes. Figure 3 (a) shows position density histograms for the charge variants. Very negative bGEMs (*−*2, 160*e*) are enriched inside and towards the periphery of the nucleoid, while the relatively neutral bGEMs (*−*840*e*) are slightly enriched outside the nucleoid but free to explore most of the bacterial cell. Very positive bGEMs (+1, 800*e*) tend to be fully excluded from the nucleoid and localize preferentially near the cellular poles. Overall, highly charged particles explore a much smaller fraction of space than their neutral counterparts, resulting in bright spots in the density histograms. In the case of the *−*2, 160*e* particles, these bright spots are often the result of motion occurring in the direction orthogonal to the projection plane shown in fig. 3 (a). E.g., little motion along *xy* and *xz* combined with considerable motion along *yz* will lead to a bright spot in the *yz* image, but not in the *xy* and *xz* images.

**Figure 3.**
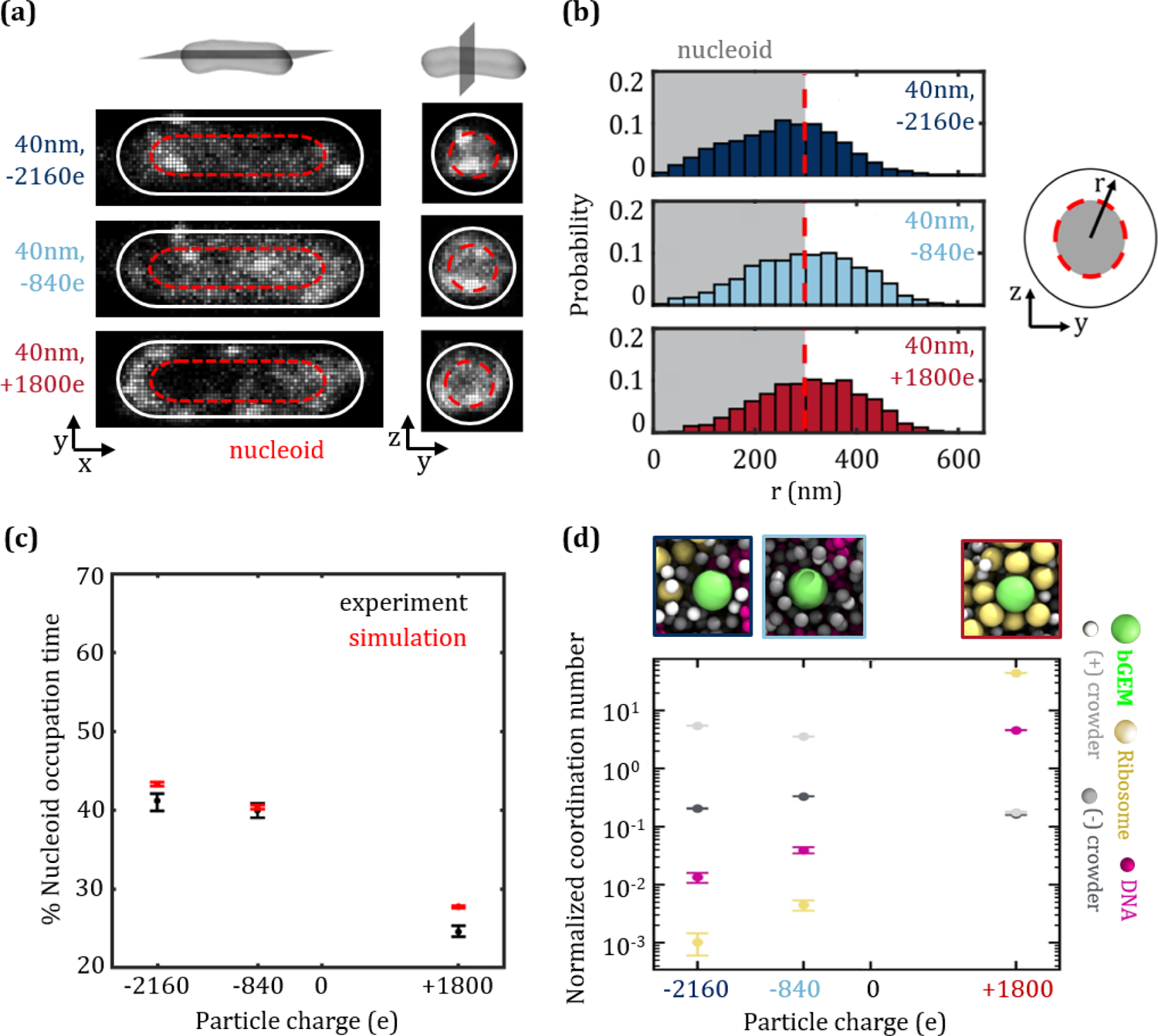
Electrostatic interactions between bGEMS and other cytoplasm macromolecules determine the localization of bGEMS. (a) *xy* and *yz* 2D histograms from experiments for 40nm bGEMs of charge of −2160e (7,500 localizations), −840e (9,500 localizations), and +1800*e* (11,000 localizations) in top, middle, and bottom images, respectively. Brighter intensity shows more localization. Contrast adjusted linearly to improve visibility. Average nucleoid region (red dashed line) and cell periphery (solid white line) are highlighted. (b) Radial density distributions (experiments) shift towards the cell periphery as net charge increases. (c) Nucleoid occupation time for experiment (black) and simulations (red) shows good agreement. Negatively charged bGEMS spend *≈*40% of their time inside the nucleoid, much more than positively charged bGEMs (*≈*25%) (d) The normalized mean coordination number (average number of neighboring negative crowders, positive crowders, nucleic acids, and ribosomes per GEM divided by that of a GEM with zero net charge) obtained from simulation quantifies macromolecular interactions and shows that positively charged bGEMS bind to ribosomes, explaining why they spend less time inside the nucleoid. Strongly negative particles form clusters with surrounding positively charged proteins, yielding a larger effective particle size yet still manage to enter the nucleoid. In contrast, only strong interactions with ribosomes outside the nucleoid and DNA inside the nucleoid definitively segregate biomolecules either inside (DNA-bound) or outside (ribosome-bound) the nucleoid. Insets: simulation snapshots for (left) −2160e, (middle) −840e, and (right) +1800e bGEMs.

In Figure 3 (b), the probability distributions shift towards the cellular periphery as charge becomes less negative. This effect is less pronounced in the radial distribution for the positive particles, however it is very clearly demonstrated by a marked increase in the polar occupation time of these particles (*≈* 25%) with respect to their negative counterparts (*≈* 15%). We next measured the amount of time the particles spent inside the nucleoid (Fig. 3 (c)). Once again there is excellent agreement between experiment (black) and simulations (red). We observed little difference in nucleoid occupation time between the two negatively charged particles, with both particle types spending close to 40% of the time in the nucleoid region. However, for positive particles, the nucleoid occupation is reduced to *≈* 25%. This nucleoid exclusion is stronger than the exclusion of the larger 50 nm particles (*≈* 30%) in fig. 2.

Our whole-cell colloidal simulations enable us to investigate the intermolecular interactions of the bGEMs with other cytoplasmic macromolecules. To quantify these interactions, we calculated coordination number probability distributions for each of the different cytoplasmic components interacting with bGEMs in the simulation. We also calculate the average coordination number for each molecule type normalized by that for a bGEM that has zero net charge (Fig. 3 (d)). The coordination number corresponds to the average number of each type of molecule whose surface is within a 1-nm shell around a bGEM. We observe that positive bGEMs are almost exclusively surrounded by ribosomes, exhibiting long-lasting interactions. Less frequently we observe additional interactions with DNA, however even in this case, the ribosomes partially surrounded the positive bGEM (Fig. S14). Ribosomes, composed primarily of negatively charged rRNA molecules, have a total charge of (*≈* −3,000 e), resulting in a stronger attraction to positive bGEMs than the nuclear DNA (Fig. S13). Further, unlike the DNA, which exists in a large continuous network, ribosomes are freely mobile and able to reorganize to surround the bGEMs. As previously discussed, active ribosomes and polysomes tend to be excluded from the nucleoid, enhancing the bGEM localization even further. This ribosome cloud surrounding the positive bGEM increases the effective particle size considerably and prevents it from moving into the DNA-rich areas of the cell.

We also find that negative bGEMs have only transient short interactions with other cellular components, resulting in much smaller coordination numbers (Fig. 3 (d),(Fig. S14 (d))). The most negatively charged, *−*2, 160*e*, bGEMs form transient clusters with positively charged crowders that are found throughout the cytoplasm and nucleoid, as evident from simulation snapshots and our calculated coordination numbers. The more neutral, *−*840*e*, bGEMs exhibit generally weak electrostatic interactions without much difference between positive and negative crowder coordination numbers. From these differences in the negative particle’s electrostatic interactions, we observe that even if large-scale localization (quantified by nucleoid occupation) is almost identical, our model suggests that bulk charge leads to significant changes in local macromolecular environments for each GEM. Molecular charge can thus have a strong impact on sub-cellular localization and organization, depending on the relative charge and localization of other cellular constituents.

### Particle dynamics are governed by charge and size

Our localization results indicate that increased macromolecular charge has an important effect on the strength and extent of interactions between cellular components. However, as exemplified in the simulation video (SI Movie 4), the bacterial cytoplasm is a highly dynamic system with constant particle rearrangements, and while stronger localization is often biologically favorable, there could be a trade-off when it comes to dynamics. To explore this, we calculated the ensemble-averaged mean squared displacement (MSD) as a function of lag time for both experimental and simulation trajectories. Figure 4(a) shows experimental MSDs for all particle sizes and charges. Focusing on the longest-time lags, we see that all the curves plateau (inset). This is expected for particles confined to a finite volume, and it demonstrates that the particles are exploring the full volume in which they are confined. In the case of the larger particles, this corresponds to the full cell size, whilst the 20nm bGEMs are mostly confined to the smaller nucleoid. This is consistent with our localization results in Figure 2. Additionally, we observe that regardless of charge or size, the motion appears sub-diffusive for all bGEMs within the first 5 timelags, i.e., with a power law exponent *α <* 1 for MSD= 6*Dt^α^*. *α* is approximately 0.75 for large particles and 0.45 for the small 20-nm particles (Table S2). These exponents were estimated by fitting the data to a sub-diffusive model that accounts for localization errors as described in §S2.

**Figure 4.**
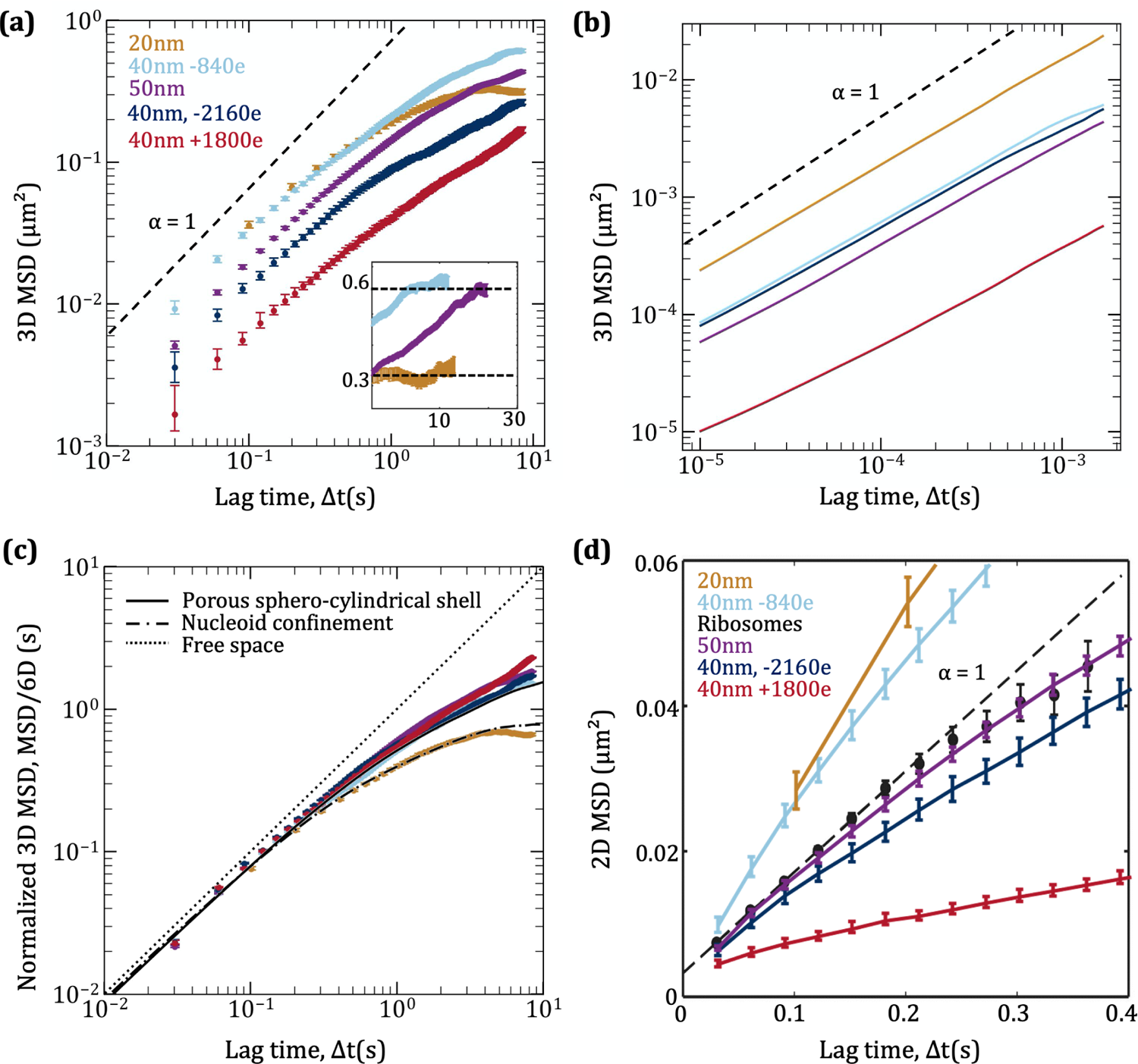
Macromolecular dynamics are governed by size and charge. (a) 3D experimental Mean Square Displacement (MSD) for all bGEM sizes and charges as a function of lag time. Motion appears sub-diffusive with an exponent smaller than 1 (dashed black line) for all particles. Inset: particles experience different confinement volumes based on size. (b) MSD from whole-cell colloidal simulations for all bGEM sizes and charges as a function of lag time. Dynamics at shorter timescales have a power-law exponent much closer to unity. (c) Experimental mean-squared displacement normalized by diffusion coefficients obtained from probabilistic simulations of the excluded volume nucleoid as a function of lag time shows that the power-law exponent of diffusion for each particle size and charge is explained by a confined random walk in the cellular region the particle occupies. (d) 2D (*x*,*y*) MSD of ribosomes and bGEMs as a function of lag time. Qualitative agreement between these data from bGEMs and previously published ribosomal diffusion data (reproduced with permission from Bakshi et al. [22]) indicates that bGEMs serve as good probes of biophysical dynamics of large particles in *E. coli*.

We next set out to explain the origin of the apparent sub-diffusivity of the bGEMs. The sub-diffusion we observe could be an effect of confined motion in a spherocylindrical cell (apparent sub-diffusion), or it could be the result of viscoelastic interactions in the extremely crowded and polydisperse cytoplasm (true sub-diffusion). Even for the smallest lag times in our 3D SPT experiments, 30 ms, there is enough time for a bGEM to diffuse a distance several times larger than its own diameter, which could introduce confinement-related artifacts into the MSD. This realization prompted us to conduct further investigations into the power-law exponent observed in experimental results using our whole-cell colloidal simulations. These simulations have a time resolution from milliseconds down to picoseconds, far below experimentally resolved timescales. In the simulated trajectories (4 (b)), we observe that departures from normal diffusion appear to be much smaller than observed in experiment (*α >* 0.84 for all bGEMs). This departures from normal diffusion is driven in part by bGEM dynamics inside the nucleoid, where motion is more subdiffusive (Fig. S15) due to particles diffusing through a stiff polymeric network. Furthermore, bGEMs outside of the nucleoid explore only a small, 383-nm, space between the cellular membrane and nucleoid, likely experiencing caging at intermediate timescales due to confinement [45]. While this caging might be enhanced by electrostatic interactions which form more durable cages and bonds, electrostatic interactions are likely not a significant source of sub-diffusion since the exponents for all bGEM charge variants are very similar (Table S2). Regardless, the dynamics observed in our simulations are driven by colloidal-scale forces and phenomena, since our model assumes that the cytoplasm is inherently Newtonian. The low amounts of sub-diffusion we find in the colloidal simulations emerge from interparticle correlations driven by confinement, crowding, and electrostatics.

Next, we further interrogated computationally the impact of confinement and geometry on long-time self-diffusion at timescales similar to those in SPT experiments (*∼* 10 ms and beyond). While a spherical enclosure is highly efficient computationally and provides accurate enclosure effects for short-time diffusion, at long times, the sphero-cylindrical shape of the enclosure matters and can influence the power-law behavior. To increase computational speed and efficiency, simulate experimental timescales, and faithfully represent cell geometry effects in the long-time limit, we expanded a probabilistic simulation method previously used in literature [22, 57, 24, 58, 36]. In these longer time simulations, the nucleoid was replaced with an excluded volume region, and a single bGEM undergoes diffusion inside of a spherocylindrical cell, with a defined probability that the GEM particle could enter or exit the nucleoid. This probability was extracted from both experimental and simulations data, and allowed rapid simulation of many trajectories. For large particles, simulating diffusion throughout the entire cell gives excellent agreement with the experimental data (Fig. S16 B-E). Meanwhile, for the 20-nm particles, the best fit corresponds to motion confined to the nucleoid exclusively (Fig. S16 C), which is consistent with the experimental radial probability distribution in Figure 2 (b). Using this model, and by adjusting the diffusion coefficients by less than a factor of 2 from those determined from the experimental MSD, we observe almost perfect agreement with the experimental data (Fig. S16) and interpret these small corrections to the diffusion coefficients as the result of imperfect fitting due to the nonlinear nature of the experimental MSD curves.

Our simulations show that 3D confinement effects are sufficient to explain the anomalous diffusion measured in *E. coli*. To further exemplify this point, we normalized the experimental MSDs for all bGEMs by the fitted anomalous diffusion coefficient (for details on the fitting refer to §S2) in Figure 4 (c). Upon doing this normalization, all the MSD curves collapse together at short and intermediate times, demonstrating that confined Brownian diffusion is the unified mechanism of apparent anomalous transport in *E. coli*. At longer times, the MSD corresponding to the 20-nm bGEMs deviates significantly from the rest, due to the faster timescales in which confinement effects begin to dominate particle motion. Overlaying the collapsed curves with the normalized MSDs from the simulations reveals that the larger particle’s dynamics are well reproduced by a particle confined to a porous spherocylindrical shell. The observed dynamics for the 20-nm particles are consistent with confinement to the nucleoid. We conclude that a large fraction of the non-trivial anomolous dynamics observed in ours and previous experiments are a result of confined diffusion to the finite volume explored by the particle, i.e. this is apparent sub-diffusion as defined above. The confinement volume is determined primarily by the size of the particle, with small contributions from particle charge (Fig. S16).

How do macromolecular size and charge affect dynamics *in vivo*? One of the features we observe is that macromolecular interactions play a key role in driving qualitative differences in dynamics over short timescales. Smaller macromolecules diffuse further than larger ones over short times, which we observe both in simulation and experiment. However, at longer time lags we observe experimentally that 20-nm particles moved approximately 50% less than naively expected as they experienced stronger confinement in the nucleoid region and increased difficulty diffusing in the porous DNA network.

Although particle charge has a more modest effect on localization than size at whole-cell length scales, it has a more appreciable impact on particle dynamics. Positive bGEMs in both experiment and simulation move considerably slower than all the other bGEM variants. While positive bGEMs are not permanently bound to ribosomes or DNA, electrostatic interactions with these cytoplasmic components slow down dynamics by almost an order of magnitude with respect to the more neutral *−*840*e* bGEMs (Fig. 4 (a) and (b)). For the very negative *−*2, 160*e* bGEMs we observe an important *≈* 40% decrease in motility with respect to the *−*840*e* bGEMs in experiment. However, in our simulations, mobility of *−*2, 160*e* bGEMs was reduced only 5-10% compared to *−*840*e* bGEMs. The smaller effect shown in simulation is a result of transient interactions with positively charged macromolecules in the cytoplasm, suggesting that an isotropic model where the net charge is used may not be enough to reproduce dynamics of very negative GEMs. *In vivo*, it is possible that GEMs have longer-lasting electrostatic bonds to positive surfaces of net negatively charged macromolecules. Other differences in experimental and simulation dynamics, such as faster dynamics observed in simulation for all GEMs, are likely due to model assumptions that were necessary to make the simulations computationally tractable, such as the absence of hydrodynamic interactions between macromolecules and between macromolecules and the membrane [43, 44, 45]. We do not make any ad-hoc adjustments to interaction parameters that may also enable a better quantitative match as done in previous studies [9]. Overall, the qualitative agreement with simulations and experiments supports our hypothesis that molecular-level interactions that act on short timescales largely set the relative dynamics over the longer timescales observed in experiments.

Given that our experimental data is best fit by a confined random-walk model in a sphero-cylindrical geometry (Fig. 4c), we converted the diffusion coefficients extracted from the long-time probabilistic simulations to an “effective viscosity” (*η*_eff_) of the cytoplasm using the Stokes-Einstein relation. All bGEMs experience a viscosity *∼* 100 cP (see Table S7), roughly an order of magnitude higher than that of proteins [59], including GFP [60, 19, 61], diffusing in the *E. coli* cytoplasm, and two orders of magnitude higher than the viscosity of water. The effective viscosity was roughly 100 cP for all particle with roughly neutral charge. However, electrostatic interactions greatly influence the effective viscosity experienced by the 40-nm charge variants, with *η*_eff,+1800*e*_ *≈* 5.5 *η*_eff_*_,−_*_840*e*_ and *η*_eff_*_,−_*_2160*e*_ *≈* 2.5 *η*_eff_*_,−_*_840*e*_.

bGEMs are useful probes for investigating the *in vivo* dynamics of similarly-sized macromolecules. To compare our results with 2D-MSD data from the literature, we obtained 2D-MSD (*x*,*y*) curves from our 3D data. In Figure 4(d), we overlay these curves for all bGEM particle sizes and charges on top of data from Bakshi et al. for ribosomal diffusion [22]. The MSD calculated from ribosome dynamics quantitatively matches our observations using 50-nm bGEMs. Like our engineered particles of this size, ribosomes (and polysomes) are highly negative [12] and tend to be excluded to the cellular periphery [22, 20]. This suggests that, as a first approximation in terms of their dynamics, ribosomes behave like spherical complexes with a homogeneous negative charge distribution on their surfaces, as reported in [12], though deviations between simulation and experiment for the reduced mobility of negative bGEMs suggest that non-isotropic interactions may quantitatively change dynamics.

## DISCUSSION

Our synergistic experimental and computational approach to studying intracellular dynamics has revealed key details of how macromolecules move in the bacterial cytoplasm. Relative speeds can be attributed to interactions within the cytoplasm whereas sub-diffusive power-law exponents observed in experiments for tracer molecules and ribosomes [34, 31, 16, 32, 16] primarily stem from three-dimensional confinement effects. These effects become evident even at the shortest experimental timescales.

First, we establish that particle size is the primary determinant of bGEM localization inside *E. coli* cells. With small particles (20 nm) enriched inside the nucleoid, medium sized particles (40 nm) able to explore both regions, but slightly enriched outside the nucleoid, and large particles preferentially excluded from the DNA-rich area (50 nm). Detailed characterization of the simulated nucleoid pore size distribution shows how size determines the percentage of nucleoid volume accessible to a given macromolecule. Our results suggest that the nucleoid acts as a size-selective migration filter, allowing smaller biomolecules to pass through easily and excluding larger complexes such as active ribosomes and polysomes. These results are consistent with a reported nucleoid mesh size of *≈* 50 nm [18] and expand on previous localization observations based on 2D-SPT experiments in the literature [18] by providing additional nuance: small macromolecules such as ribosomal subunits are able to freely diffuse into the bacterial nucleoid, and are likely to be enriched in this region. Large macromolecules, on the other hand, tend to be preferentially, but not *completely*, excluded from the nucleoid region, which is exemplified by the 50-nm bGEMs spending roughly 30% of their time inside this region.

These results suggests that size-based localization could serve as an important mechanism for biophysical regulation. A particularly striking example of the biological importance of size-based filtering is the mRNA life cycle, and the spatial segregation of transcription and translation inside the cell. It has been proposed that mRNA molecules are transcribed from DNA inside the nucleoid, where ribosomal subunits assemble around the molecule to start translation. The ribosomal-RNA complex is then exported towards the cellular periphery [62], and finally after translation the RNAs are degraded by RNAses located near the inner membrane of the cell [63]. The mechanism for these processes is not fully understood, but our observations suggest that small ribosomal subunits and inactive ribosomes (*≈* 20 nm) might be preferentially enriched inside the nucleoid, where the nascent mRNAs are found. Once the translation process begins, and as the mRNA-ribosomal complex grows in size, its probability of localizing towards the periphery increases, and thus the complexes migrate outside the nucleoid. Hence, once the mRNAs are fully translated, they will likely be at the correct location for degradation. Future work will be needed to explore how the nucleoid microstructure might be tuned by changing solvent or growth conditions to promote or hinder the migration of different biomolecules such as transcription factors, ribosomes, RNA polymerase clusters.

We report that the localization of particles shifts towards the cellular periphery as charge becomes less negative. Positively charged bGEMs (+1, 800*e*) are almost completely excluded from the DNA-rich area, localizing preferentially at the cellular poles. Furthermore, although negatively charged bGEMs occupy the nucleoid for similar amounts of time regardless of absolute charge (*−*2, 160*e* versus*−*840*e*), positive bGEMs spend a much smaller fraction of time inside the nucleoid than any other bGEM across all sizes and charges. Previous studies in *E. coli* inferred abundant interactions between positive proteins and ribosomes [39]. Similarly, our dynamic simulations suggest that positive bGEMs are surrounded by highly motile ribosomes and polysomes in the cytoplasm, which are themselves entropically segregated from the nucleoid. Our findings suggest that sub-cellular particle localization is an emergent property of multiple underlying phenomena. In this case, localization is primarily due to the coupled entropic and electrostatic effects of interacting, polydisperse macromolecules diffusing in a porous medium and confined to a finite space. Our simulation results also show that, at a smaller length scale, the neighboring environment of macromolecules is dependent on macromolecular charge, which suggests that electrostatic interactions might drive increased localization of pairs of molecules that can speed up search processes within the cell, as proposed previously [26, 27].

Although increased intermolecular interactions might be beneficial for particle localization, the strength of these interactions can lead to decreased motility in the cell [41, 27]. Indeed, we observed this trade-off in our experimental and simulation results, where increased net charge of the bGEMs increases electrostatic interactions leading to reduced apparent diffusion coefficients. In particular, positively charged 40 nm bGEMs effectively diffuse as much larger particles due to interactions with ribosomes, often forming large clusters. This slowdown of dynamics and the overall difficulty to access the nucleoid region, might explain in part the charge/size hierarchy in bacterial cells and the absence of positive large macromolecules. While slower dynamics for positively charged particles has previously been reported in the literature for bacterial and eukaryotic proteins [39, 64], in both of these studies, significant differences in dynamics were not observed for negatively charged macromolecules of small and large net charges. In our experiments, we observe a large difference in the effective diffusion coefficient between the two 40-nm negative bGEM species. This trade-off between localization and dynamics may be tuned in the cell to avoid dynamic arrest and ensure small enough times for search and capture processes that require physical proximity of the cellular components [2].

Although our results reveal that the nature of the diffusive process inside bacteria is simple diffusion in a crowded environment, we see a rich array of dynamic behaviors at different timescales. In particular, at the timescales of many biological processes, macromolecules effectively experience sub-diffusive motion due to confinement. By tuning properties such as a molecule’s charge and size, cells may be able to achieve localization to a particular region of the cell. This localization might improve the odds of specific reactions occurring and decrease search times, freeing up energetic resources for other processes in the cell.

## METHODS AND MATERIALS

### Experimental setup

#### Bacterial Genetically Encoded Multimeric nanoparticles (bGEMs)

In this work we engineered 5 new bacterial strains summarized in Table 1. The bGEMs are expressed through new plasmids developed for this study (see S0 in SI for details), primers and strains used in this study can be found in tables S3, and S4. Plasmids were transformed into MG1655 WT *E.coli* using TSS transformation. We based our probe design on the original Genetically Encoded Multimeric Nanoparticles (GEMs) developed by the Holt lab for tracking experiments in eukaryotic cells [55]. GEMs self-assemble into bright monodispersed particles with a defined size and shape from homomultimeric scaffolds fused to a fluorescent protein [55] (Fig. S4. a). Prior to this work, two types of GEMs were published in the literature [55], with 20 and 40 nm diameters. For this study we developed a new 50 nm GEM particle, measured by negative stain EM (r=25 *±* 0.5 nm), using genes from the encapsulin protein of the hyperthermophilic archaeon *Vulcanisaeta distributa* [65] (see Fig. S4. b, and additional details in S0 in SI). Additional to the bGEMs plasmids, we used a second plasmid (pSACT101 Amp*^r^*) to label the DNA using the nucleoid associated protein HU-alpha labeled with mRFP1.

**Table 1.** bGEMs strains developed.

| Strain number | bGEM size (nm) | Fluorescent protein | Antibiotic Resistance |
| --- | --- | --- | --- |
| DVM85 | 20 | msfGFP | Kan<sup>r</sup>, Amp<sup>r</sup> |
| DVM76 | 40 | msfGFP | Kan<sup>r</sup>, Amp<sup>r</sup> |
| DVM79 | 40 | GFP +15 | Kan<sup>r</sup>, Amp<sup>r</sup> |
| DVM83 | 40 | GFP -18 | Kan<sup>r</sup>, Amp<sup>r</sup> |
| DVM90 | 50 | msfGFP | Kan<sup>r</sup>, Amp<sup>r</sup> |

#### Charge variants experimental design

Because most of the charge will be screened by the ion rich cytoplasm, charge interactions occur locally involving only a few GFP monomers, therefore to compare particle size effects, we used the msfGFP’s native charge of *−*7*e*. To explore charge effects, we used the 40 nm diameter scaffold fused two 2 additional GFP charge variants, +15*e* [66] and *−*18*e* (developed for this study, see §S0 in SI and Fig. S4.c) resulting in three bGEM charges with total particle charge: *−*2160*, −*840 and +1800*e*.

#### Bacterial sample preparation

All cells were grown in EZ-rich media [67]. For the stationary phase experiments, cells were grown from frozen in the shaker at 37 *^◦^*C for roughly 20 hours. Then 100 *µ*L aliquots were prepared with 1 *µ*L of 0.5 mg/*µ*L WGA 405(S) cell-surface fluorescent label and incubated for an additional 30 minutes to an hour shaking at 37 *^◦^*C. For stationary phase experiments the use of induction was not required to achieve the desired bGEM assembly. For exponential phase experiments cells were grown overnight until stationary phase. Then, the cultures were back-diluted 1:100 in EZ-rich media and grown in the shaker for 1 hour at 37 *^◦^*C. This short incubation time was enough to reach early exponential phase but still preserve the bGEMs that formed during the long overnight culture. At this point 100 *µ*L aliquots were prepared with 1 *µ*L of 0.5 mg/*µ*L WGA 405(S) membrane stain. For the 20nm bGEMs 1 *µ*L of 100 ng/mL anhydrous tetracycline was added for bGEM induction. The aliquots were then incubated for an additional 30 minutes shaking at 37 *^◦^*C. For all imaging experiments, 1.5% agarose pads were prepared with the corresponding nutrient conditions. For exponentially growing cells pads were made with EZ-rich media.

In the case of stationary phase cells, we used M9 media without a carbon source. Pads were made using a tunnel slide. For all growth conditions, 1 *µ*L of cells was placed in each imaging pad and allowed to dry. The samples were then covered with a coverslip and sealed with Valap (1:1:1 mixture of vaseline, lanolin and paraffin).

#### Imaging

All single particle tracking microscopy was performed with a custom built biplane wide-field illumination microscope (Fig. S5 for optical diagram). Images were obtained using a 1.49NA 100x Nikon objective and 2x optical magnification, and captured with an Andor EM-CCD camera. The setup includes bright field illumination and 3 colors for epifluorescence: 405 nm, 488 nm and 561 nm. To enable 3D tracking, we use a biplane module consisting of a mirror and beam-splitter (see §S1 in SI for more details). The optical distance between the two focal planes is *≈* 525 nm. All imaging was performed at room temperature (23.3 *^◦^C*), controlled to 0.5 *^◦^*C. Samples were imaged from the bottom through a coverslip, using oil matched to the diffraction index of glass. An exposure time of 30 ms was used for all samples except for the 20-nm bGEMs, which required an exposure time of 100 ms due to decreased particle brightness. Each datapoint in the experiments consists of 3 channels. A red channel with a nucleoid *z*-stack, a blue channel with a membrane *z*-stack and a green channel movie with 500-1000 frames of bGEMs motion inside the cell (SI Movie 2). All *z*-stacks consist of 40 steps spaced by 100 nm, with the stack center corresponding to equal focus on both biplane images. A sample data set with 3 timeframes is shown in Figure 1. Notice how each image seems duplicated at the top and bottom. This corresponds to the two biplane images produced in the same camera chip. All images were analyzed to determine the *xyz* positions of the particle at each timeframe using a custom built image analysis pipeline in Matlab[68]. Further details of the optical setup and image processing can be found in §S1 in SI.

### Negative stain transmission electron microscopy for 50nm GEMs

50 nm GEMs were purified as described in [55]. Purified particles were deposited on carbon-coated 400 mesh copper/rhodium grids (Ted Pella Inc., Redding, CA), stained with 1% aqueous uranyl acetate, examined in a Philips CM-12 electron microscope and photographed with a Gatan (4k x2.7k) digital camera.

### MSD calculations and corrections

Ensemble-average MSDs were calculated using custom code developed in Matlab and Python. Experimental static and dynamic localization errors, lead to artifacts in the shape of experimental MSD curves in the log-log scale. The static localization error, has a pronounced effect in our unprocessed MSDs, causing the curves to appear highly sub-diffusive for early time points when plotted in a log-log plot (Fig. S10). To correct for these errors we fitted our MSD data with a model that accounted for both types of errors [69], and subtracted the static localization error from the MSD data to obtain the plots in fig. 4. Additional details are descibed in §S2 in SI.

### Simulations

#### Whole-cell colloidal simulations

We constructed a coarse-grained whole-cell model of E. coli (Fig. 1(c)) using the massively-parallelized molecular dynamics package LAMMPS [70]. Each simulation included a spherical confining cell membrane and approximately 30,000 spherical particles representing a porous nucleoid and a polydisperse cytoplasmic milieu of positive and negative native cytoplasmic crowders (to capture crowding effects from proteins, tRNA, transcription factors, ribosomal subunits, etc.), ribosomes, polysomes, and GEMs.

We chose the radii of the nucleoid region to be 200 nm to reflect the average values measured in our experiments in *E. coli*. The nucleoid was constructed via a random self-assembly process, forming an interconnected network of polydisperse DNA beads with average radius *a*_DNA,i_ = 7nm. The self-assembly process can be tuned to produce a network structure that recovers the nucleoid morphology measured in experiments. For more detail, please see §S4 in SI and Figure S11. The radius of the cell was chosen to be 383 nm to match the relative nucleoid to cell volume of 0.39. Macromolecular abundances were determined to represent physiological data of stationary phase *E. coli* in rich growth media [51, 52, 53, 41, 54] (see Table S5) and the resulting volume fraction of the cell was *φ* = 0.31. The position of macromolecules inside the cell membrane and interpenetrating or surrounding the nucleoid was initialized in Packmol [71].

In our model, macromolecular interactions were represented with a hard-sphere [72] Debye-Hückel [42] potential, where each pair of molecules experiences entropic exclusion and screened electrostatic attractions or repulsions that depend on their relative charges (Fig. S13). These coarse-grained interaction potentials were previously derived from and parameterized by all-atom simulations [42]. In addition to these pairwise forces, DNA beads and ribosomes that are a part of the same polysome interact via a harmonic bond potential. Confinement is enforced via a harmonic force that acts in the radial direction if a particle attempts to move outside the enclosure.

Following equilibration, we ran dynamic simulations and measured the MSD and nucleoid occupation time of GEMs. The timestep was chosen to be 42 ps to maintain inertia-free dynamics. Simulations were evolved for 3.4 ms, with particle trajectories output every 0.8 *µ*s. Visualizations of the model were created using Visual Molecular Dynamics (VMD, [73]). For more details with regard to the colloidal whole-cell model, please see §S4 and §S5 in the SI.

#### Probabilistic simulation of an excluded volume nucleoid

In this type of simulation, particles are randomly instantiated in different regions of the cell. The particles perform a random walk drawing displacements from a Gaussian distribution as permitted by boundary conditions which enforce confinement within the spherocylindrical cell and entry/exit probabilities into and out of the nucleoid. Simulation parameters including cell and nucleoid shape and size, diffusivity, and nucleoid occupation fractions were obtained from experiment-based measurements and outputs from the whole-cell colloidal model. See §S6 in SI for additional details.

## AUTHOR CONTRIBUTIONS

D.V-M, A.M.S, B.P.B, Z.G., R.N.Z, and J.W.S designed research; D.V-M performed bacterial experimental work and developed image analysis pipeline; A.M.S and J.L.H developed colloidal simulations; A.M.S and D.V-M developed MC simulations; D.V-M and J.P.S developed bacterial strains; L.J.H and M.D developed the 50nm GEM particle; D.V-M, A.M.S, B.P.B performed data visualization and analysis; D.V-M, A.M.S, R.N.Z, and J.W.S wrote the manuscript

## ACKNOWLEDGMENTS

This work was supported by the NSF, through the Center for the Physics of Biological Function (PHY-1734030) and GRF 1656518 to A.M.S and J.L.H; and a Joseph H. Taylor Fellowship to D.V-M. Crispin Stanford Graduate Fellowship to A.M.S, and an ARCS Foundation Award for J.L.H; L.J.H. was funded by NIH R01 GM132447, R37 CA240765, the NIH Director’s Transformative Research Award TR01 NS127186, and the Human Frontier Science Program (RGP0016/2022-102). Computing was performed on the Sherlock cluster at the Stanford Research Computing Center (SRCC) and on Stampede2 at the Texas Advanced Computing Center (TACC). SRCC resources were provided by Stanford University, while resources at TACC were supported by the National Science Foundation’s Extreme Science and Engineering Discovery Environment (XSEDE) Research Award No. CTS120035 and Advanced Cyberinfrastructure Coordination Ecosystem: Services & Support (ACCESS) Research Award No. CHM230008. We thank David Liu for providing the genetic material for the GFP charge variants, Nick R. Martin for providing genetic material for plasmids, Dajun Sang for help with GEM purification and EM, Matthew E. Black for insightful discussions regarding experimental details, Brian K. Ryu for help with void-width calculations, and Theo S. Yang for suggestions regarding modeling nucleoid polymer dynamics.

## Supporting Information for

### Supporting Information Text

#### S0. bacterial Genetically Encoded Multimeric nanoparticles

##### bGEMs characteristics and design

GEMs self-assemble inside the cell from multiple sub-units coming together in fixed stoichiometries to form monodispersed icosahedral particles [1]. These sub-units are protein complexes consisting of a protein scaffold and a fluorescent protein (FP) linked together (Fig. S4.A). The scaffold protein determines the size of the assembly, and the resulting particle is fully coated by the fluorescent proteins. Thus all interactions between the GEM and the environment occur through the FPs. The scaffold proteins originate from hyperthermophilic archaea and bacteria that are vastly different from *E. coli* minimizing the risk of biological interactions. To develop the new 50 nm GEM we used scaffolding domains based on the encapsulin protein from the hyperthermophilic archaea Vulcanisaeta distributa [2]. Using negative stain electron microscopy, we measured a radius of 25.03 ± 0.52 nm. (Fig. S4.B). The 50 nm particles were originally conceived for research in eukaryotes, and were synthetized following the steps described in [1]. In brief, the Vuldi encapsulin was synthesised with an IDT gene block and codon-optimized for eukaryotic expression. Gibson cloning was used to introduce the encapsulin into a yeast expression plasmid after the INO4 promoter. A Sapphire fluorescent protein was fused at the C-terminus.

Particles are expressed through a low copy-number plasmid with an anhydrous tetracyclin (aTet) promoter. We replaced the original t-sapphire fluorophore in [1] with msfGFP for all three size variants (20, 40 and 50 nm). For the 40nm particles we used 2 additional fluorophores to vary the total charge of the GEM, GFP +15 from [3] and a new GFP-18 flurophore developed here. We sequenced every charge variant to verify the charge of the constructs. We determined the charges from the sequence data using ProteinCalculator V3.4 (https://protcalc.sourceforge.net/). The size variants are all coated with standard msfGFP of charge −7e, making them close to neutral in terms of their electrostatic interactions. The total charge of a particle corresponds to the charge of the individual GFP multiplied by the number of subunits making up the particle, e.g. for the 40 nm “neutral” particle the total charge is −7e·120 = −840e. Thus, for our charge analysis we used three charge variants of the 40 nm sized particles: −2160e (GFP −18e), −840e (msfGFP −7e) and +1800e (GFP +15).

##### Strains construction

All the plasmids developed for this study were constructed using the Gibson Assembly [4] kit (New England Biolabs, MA). For the plasmid pDVM1, containing PFV-msfGFP, the PFV gene (40 nm particle scaffold) was amplified from pLHC611-pcDNA3.1-Pfv-Saphire [1] using primers DVMO3 and DVMO22, the msfGFP gene was amplified from NMP158 [5] using primers DVMO20 and DVMO21. The two genes were fused in frame and inserted into the pEVS143 [6] backbone, an expression vector containing an anhydrous tetracycline promoter and kanamycin resistance cassette. pDVM1 was used as the backbone to construct the other plasmids used in this study. We used primers DVMO37 and DVMO38 to amplify the backbone around the PFV gene to replace the scaffold and the primers DVMO43 and DVMO44 to amplify the backbone around the msfGFP gene to replace the fluorophore. For the plasmid pDVM16, containing AqLS-msfGFP, the AqLS gene (20 nm particle scaffold) was amplified from pLH1426-pRS305-PIN04-AqLS [1] using primers DVMO39 and DVMO40, and inserted into the pDVM1 backbone, replacing the PFV gene. For the plasmid pDVM26, containing Vuldi-msfGFP, the Vuldi gene (50 nm particle scaffold) was amplified from pLH1426-pRS305-PIN04-Vuldi (this study) using primers DVMO41 and DVMO42, and inserted into the pDVM1 backbone, replacing the PFV gene. For the plasmid pDVM30, containing PFV-GFPPOS15, the GFPPOS15 gene was amplified from pET-GFP-POS15 [3] using primers DVMO45 and DVMO46 and inserted into the pDVM1 backbone, replacing the msfGFP fluorophore. For the plasmid pDVM34, containing PFV-GFPNEG18, the GFPNEG30 gene was amplified from pET-GFP-NEG30 [3] using primers DVMO45 and DVMO46 and inserted into the pDVM1 backbone inserted into the pDVM1 backbone, replacing the msfGFP fluorophore. Upon sequencing a big sequence mutation in the GFP’s sequence was discovered, with the charge of the actual fluorophore corresponding to −18 e instead of −30 e. The mutation occured during cloning as the GFPNEG30 sequence was verified by sequencing. See Figure S4.C for full GFPNEG18 amino acid sequence. Fluorescence of the new fluorophore was adequate and bGEM particles assembled correctly without clumping.

The plasmids were electroporated into S17 cells and selected on kanamycin plates, the resulting plasmids were verified by sequencing and then transformed into MG1655 using TSS transformation. Adequate particle formation was checked by fluorescence microscopy. Additional to the bGEMs plasmids, a second plasmid (pSACT101 Amp*^r^*) expressing a protein fusion of the nucleoid associated protein (NAP) HU-alpha labeled with mRFP1 was transformed into the cells using TSS transformation and verified for resistance to ampicillin. Fluorescence microscopy was used to verify correct nucleoid labelling, with a Vybrant DyeCycle Violet Stain [7] (Invitrogen) as a control. Tables S3 and S4 summarize the primers and strains used in this study.

#### S1. 3D single particle tracking using biplane microscopy

Single particle tracking (SPT) consists of successive detection and localization of a particle over a sequence of timeframes. These particle positions are then stitched together to form trajectories [8]. In this work we used biplane microscopy for 3D-SPT, here we provide details regarding the experimental setup and the image analysis pipeline.

##### Custom built biplane microscope

All imaging was performed with a custom built biplane wide-field illumination microscope. A simplified optical diagram of the microscope is shown in Figure S5. To enable 3D tracking, we use a biplane module consisting of a mirror and beam-splitter. The module is placed just before the camera, as shown in Figure S5. When the light from the sample reaches the beam splitter, 50% of the light travels straight through the beam splitter and focuses in the bottom half of the camera (bottom plane), while the other 50% of the light is diverted by the beam splitter, reflecting off the mirror and into the top half of the camera (top plane). Thus the light deflected by the beam splitter travels a longer optical path length than the light that continues straight. This results in two images of the same object at different focal planes obtained simultaneously in the same camera chip. The optical distance between the two focal planes is ≈525 nm. In the following sections we’ll refer to the biplane images as planes.

##### Experimental PSF determination

The experimental measurement and processing of the PSF was obtained following a procedure similar to those described in [9, 10]. In brief, we imaged 100 *z*-stacks of 40 nm carboxylated beads using the biplane setup, with a range of ±2*µm* around the best overall focus for both planes, resulting in thousands of images of the PSF. We applied a bandpass filter, to remove fast and slow varying noise from the images. Then the coordinates corresponding to the center of each bead in *xy* were identified for both top and bottom planes. Two sets of *z*-stacks (top and bottom) were extracted for each bead using a 9×9 box around the *xy* position of the bead in each plane. The resulting stacks were summed for all beads to obtain a top and bottom calibration PSF. Gaussian filters were then applied to the calibration stacks with *σ* = 1 pixel in *xy* and *σ* = 2 pixels in *it*. Finally, the resulting top and bottom PSFs were normalized.

##### Image analysis pipeline

All software used for image analysis was custom developed in Matlab [11] and is available in GitHub (PrincetonUniversity/shae-biplaneMicroscopy-public). The input of the pipeline consists of the membrane, nucleoid and GEM channels. For the membrane and nucleoid we selected an average of the stack slices with the best focus on the bottom plane. The nucleoid and membrane channels are used for cell segmentation and background subtraction, while the GEM channel is used for tracking. A sample data set is shown in figure 1 of the main body of the paper.

To map the images corresponding to the two planes, we use a thin plate spline (TPS) [12]. Once this mapping was found, we added the two sub-images together to create a composite image from the two separate focal planes (S6.A). This composite image was used for the *xy* peak detection. Before peak detection two important pre-processing stages were performed: cell segmentation and background subtraction. (Figures S6.B and S6.C).

##### Cell segmentation and background subtraction

Because of the high density of cells in a typical image, cell-segmentation was important for analyzing individual cells. For this task we use a multi-stage approach, consisting of a user based manual segmentation followed by an automated fitting algorithm. In the manual stage, the user selects cells containing GEMs by drawing a 4-sided polygon around the desired cell using a custom built GUI in Matlab. Then, the list of manually segmented cells and their coordinates are fed to an automatic segmentation algorithm. The algorithm iterates over all the cells selected by the user. First, it generates a mask from the user’s coordinates (user’s mask). Next, in the case of stationary cells, the nucleoid image is masked to keep only the nucleoid of the cell of interest. Nucleoids are then segmented using edge detection (Fig. S11.A) and dilated by 2 pixels to create and approximate cell outline. This rough outline is then characterized by measuring the cell’s approximate size, orientation and center location. These parameters are used as a starting condition for cell fitting. Figure S7 shows an example of the fitting pipeline. In the cell fitting stage the composite image is cropped around the cell of interest using the user’s mask (S7.A). Using the parameters found from the nucleoid characterization, the cropped image is fitted with a “blurry” rectangular mask using least squares. The rectangular mask is blurred by a *σ* = 1 pixel 2D-Gaussian to simulate the effects of diffraction as shown in S7.B. Then, a sphero-cylindrical mask is constructed based on the fitted parameters. Figure S7.C shows the binary sphero-cylindrical mask convolved with a *σ* = 1 pixel Gaussian filter.

For exponentially growing cells, instead of using the nucleoid as described above, the user draws a mask using the membrane images and the fitting is performed on this mask as described above.

One of the major challenges encountered during peak detection was the presence of a strong background signal coming from free (un-assembled) GFP-scaffold sub-units diffusing in the bacterial cells. The best way of improving the SNR in this scenario is to remove the background to enhance the signal from the particles. We assume a uniform distribution of fluorophores in the spherocylinder and use this knowledge to subtract the intensity of these particles from the image. In the case of the *xy* peak finding the background subtraction is simple. In brief, the fitted sphero-cylindrical binary mask is convolved with a gaussian kernel of *σ* = 1 to simulate a uniform background blurred by diffraction (*mask_sp−cyl_*) and the fitted cell is subtracted from the cropped image GEM channel (*I_min_*) according to equation 1,

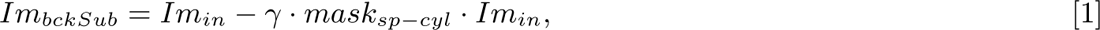

where *γ* ∈ [0, 1]. For our images *γ* = 0.6 yields the best results. The result is a background subtracted cell ready for *xy* peak detection (S7.D).

Figure S8 shows localization error histograms for simulated images with and without implementing the background subtraction algorithm. There is a considerable decrease in the localization error for the background subtracted images. The improvement in localization error is more significant in the *y*-axis because the background noise biases particle localization towards the center of the cell. Implementing background subtraction before the peak finding stage gives localization uncertainties of *σ_x_* ≈ 57 nm and *σ_y_* ≈ 68 nm for simulated images with high noise, similar to the noisiest imaging conditions we observed in our data. These errors are consistent with experimental localization errors (Table S2).

For the *z* background subtraction, the algorithm is more complex, and requires prior peak detection in the *xy* plane as described below. For the first five frames in the movie we execute the following algorithm. First, rotate the cell to align the long axis with the *x*-axis. Then normalize the two biplane sub-images by the max intensity in the movie for each plane. Then convolve the top and bottom planes of the image with their respective PSFs. We will call the resulting images from these convolutions *cBotIm* and *cT opIm*. Next we create a composite image for *z* background subtraction by doing the following weighted sum of the two image planes,

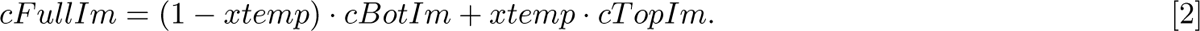

The weighing factor *xtemp* depends on the intensity ratio between the top and bottom planes for the most likely candidate particle found in the frame during the *xy* peak finding stage (*xInt* = *intT opPeak/intBotP eak*),

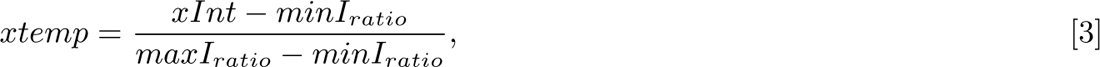

with *minI_ratio_* and *maxI_ratio_* corresponding to the minimum and maximum intensity ratios calculated from all the candidate particles in the movie.

Once we obtain this weighted composite image we then re-slice along the *y*-coordinate of the top candidate particle in the frame. This *xz* re-slice is fitted using least squares to find the optimal background subtraction, given the sphero-cylindrical nature of the cell. For the fitting step we create a template image by approximating the re-sliced sphero-cylinder to a rectangle. We place 2 rectangles centered at the focus of each biplane plane. Then the rectangles are convolved with Gaussian filters to match the convolved data. The fitting optimizes the size of the rectangles and the center of each rectangle, the intensity of the rectangles and the width of the convolved Gaussian. Once all five frames are done, the resulting fitted background images are averaged to obtain the fitted top and bottom background in the *xz* plane. These images are subtracted from the top and bottom planes during *z*-peak finding to improve localization accuracy.

##### *xyz* particle localization

For *xy* particle localization we used a 2 stage peak finding algorithm. In the first stage, we detected the brightest pixel in a window size of 15 pixels. A list of the peaks is then used in the second stage, where an intensity Gaussian is fitted to each of the peaks to find the center of the particle with sub-pixel resolution. When the fit fails, the algorithm returns the center of mass of the peak instead.

Once candidate particles are found using *xy* peak detection, we are able to determine the particle’s position along the *z*-axis. For this purpose we developed a new localization algorithm. This algorithm is robust to differences in SNR between different bacterial strains, and performed better than other algorithms, such as calibration curves, for our noisy data set.

In our approach, each plane’s image is convolved with its respective PSF to obtain a 3D stack for each timeframe. These stacks are then reshaped by slicing through the *y*-coordinate of the candidate particle to obtain a *xz* projection of the data-stack at the plane of best focus in *y*. Background subtraction (as described above) is performed on these *xz* projections to reduce background noise and entered at the *x*-coordinate of each candidate peak. We then extract *z*-slivers (10×1 pixels) of data for both top and bottom planes. Next, we fit a 1D-Gaussian distribution to the intensity profile of these slivers to find the sub-pixel localization of the particle in each plane. We then implemented a weighted sum to add these coordinates together. To determine the weighing factor we used a modified version of the Vollath 5 autofocus algorithm [13],

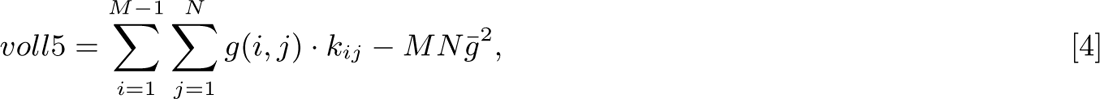

where *g* corresponds to the intensity matrix of the peak (pixels) of *M* rows and *N* columns. *k_ij_* is given by

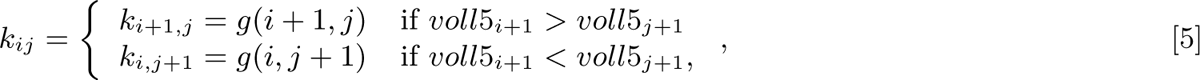

where *voll*5*_i_*_+1_ and *voll*5*_j_*_+1_ are the result of calculating *voll*5 with *k_i_*_+1,*j*_ or *k_i,j_*_+1_ respectively. The calibration parameter *c.p.* is given by the log-ratio of the Vollath 5 function applied to both biplane images,

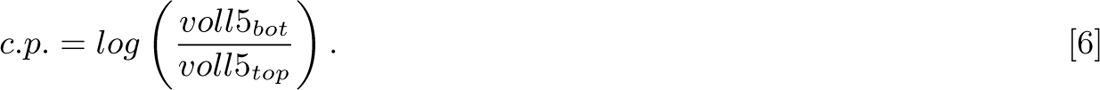

The Vollath 5 autofocus algorithm is ideal to determine the particle’s *z* position due to it’s high sensitivity to changes in position and it’s large operating range. However, it is also sensitive to the background noise of the image. Therefore the weighing parameter used to sum the particle’s locations from each plane had to account for the SNR of the candidate particle throughout the movie. This consideration is important because each type of GEM has its own intensity and noise profile so that the expected SNR range for each data set varies considerably.

The SNR was incorporated as follows. If we denote the particle’s localization in each of the planes as *z_top_* and *z_bot_*, then the particle’s *z*-position, *posZ*, is given by the weighted sum,

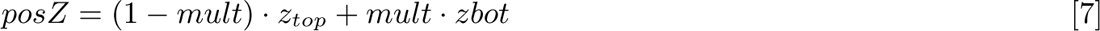

as exemplified in fig S9. The weighing parameter *mult* is given by the linear equation,

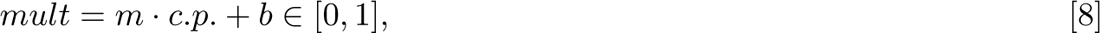

with slope 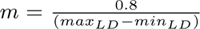 and intercept *b* = 0.9 − *m* · *max_LD_*. *max_LD_* and *min_LD_* correspond to the maximum and minimum values emxapxeLcDte−dmfionrLDthe c.p. given the SNR of the movie.

To determine the expected range for *c.p.* for a specific GEM given a movie’s SNR, we calculated the maximum SNR and the *c.p.* range [*min_LD_, max_LD_*] for both planes for all the trajectories in the data-set, and empirically determined the following relationships:

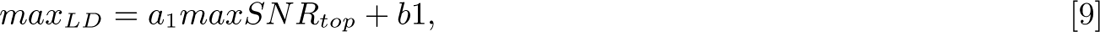

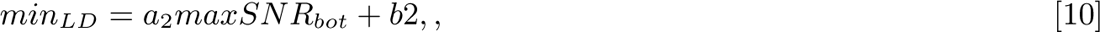

where the fitting parameters *a*_1_, *a*_2_, *b*_1_ and *b*_2_ were determined by applying linear fits to *max_LD_* = *max_LD_* (*maxSN R_top_*) and *min_LD_* = *min_LD_*(*maxSN R_bot_*). Using equations 9 and 10, we are finally able to determine the range [*min_LD_, maxLD*] for a given particle trajectory by calculating the SNR for each plane. The maximum SNR is calculated as *maxSN R* = *^maxInt^*, where *avgInt* corresponds to the mean pixel intensity in the bacterial cell and *maxInt* is the maximum intensity of the particle throughout the movie. The intensity of the particle is defined as the sum of all the pixel intensities in the window. A graphical representation of the convolution tracking algorithm is shown in Figure S9.

##### Trajectory stitching

Once the peak finding step is concluded it is time to stitch the trajectories together. To simplify tracking and stitching, we chose to work with cells that had only one GEM. This allowed us to write custom code that takes into account this information to build a trajectory throughout every frame in the movie. We developed two versions of the stitching algorithm, and for each cell we selected which of the two algorithms performed better by visual inspection. The first algorithm is a simple distance minimization. For each frame, candidate particles are tested to see which one will result in the smallest overall displacement based on an Euclidean distance metric (previous frame to current + current frame to next), if the overall displacement for a particular frame is too large, a NaN value is returned for that frame. Thus a vector is stitched for each possible starting position (if there is more than one) and the vector with the smallest trajectory length is chosen as the particle. Sometimes when using this algorithm the particle gets lost and the resulting trajectory is wrong from a certain point on-wards. To solve this problem, we developed a second algorithm. The overall functionality is very similar, however the main difference is that likely particles are found first, by selecting the 20 % brightest candidate peaks corrected for bleaching, and a trajectory vector is seeded with these probable particles. Then, frame by frame distance is minimized, using the seeded points as “anchoring” points. Depending on the noise conditions of the image, usually one of the two algorithms will perform better. Once we have preliminary trajectories (constructed with either algorithm), we interpolate to fill in any gaps. After the trajectories are finalized we do a manual correction of trajectories through a custom built GUI to eliminate any miss-localizations and bad stitches. The resulting trajectories are symmetrical in the *yz* plane and appear unbiased. Finally, all trajectories were visually inspected to discard any bad trajectories (asymmetric, with lines of data along edges, etc) resulting from experimental errors, such as imaging out of focus or bad peak localizations.

##### Intensity outliers removal

After the trajectories are stitched together we plot the intensities of the GEMs as a function of *z*-position to remove any outliers. These outliers usually correspond to out-of-focus movies or too-bright particles that are likely clumps formed by several GEMs. The GEM intensity is determined by approximating the PSF in the *z*-axis as an intensity Gaussian with amplitude *A*_0_ given by

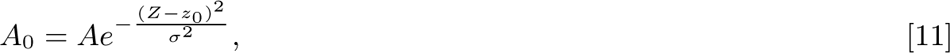

where *A* is the total measured intensity of the peak at a particular *Z*-coordinate of the PSF. The standard deviation *σ* and offset *z*_0_ are fitted from an experimental PSF obtained from 50-nm bead *z*-stacks. Using these equations, we obtain two approximations for the GEMS intensity from the two PSF intensities (*A_±_*) sampled by the biplane. With 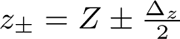, where Δ*_z_* is the optical distance (≈ 525 nm) between the two image planes.

#### S2. MSD corrections and fitting

Consider the general expression for the MSD of a diffusive process,

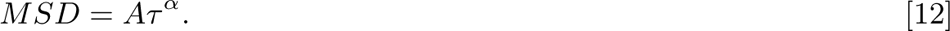

This expression is idealized assuming no measurement errors. However, in any single particle tracking experiment, the measurements will be subject to 2 main types of localization error. A static localization error and a dynamic localization error [14].

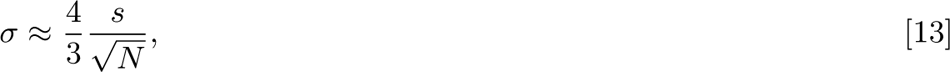

The static localization error, *σ*, arises from errors in estimating the center of the PSF and depends on the number of photons available to localize the particle. It is approximately given by [14], where *N* is the number of photons per pixel and *s* is the standard deviation of the PSF. The static localization error increases with noise.

The static localization uncertainty adds a factor of 2*σ* to the experimental MSD as shown below,

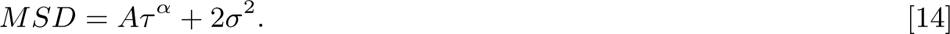

When plotting the MSD in a linear scale this error reflects in a shift of the total curve upwards, however if the MSD is plotted in a log-log scale, the resulting curve appears to be sub-diffusive for the smaller time lags [15]. This effect is clearly visible in Figure S10.

The second type of localization error is dynamic localization error, which can be thought of as a blurring error resulting from imaging for a finite exposure time *t_E_*. During the time the camera is acquiring photons, the particle is moving and thus this results in the motion of the particles appearing to be ballistic or super-diffusive for the earlier time lags [16]. Unfortunately, the effect on the MSD is considerably more complicated than the effect of the static error as described by Savin and Doyle [16],

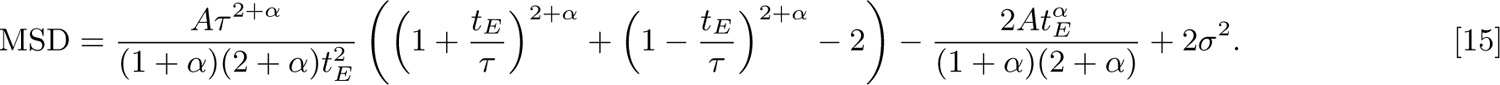

The last term in this equation corresponds to the static localization error.

For our MSD analysis we fit our curves to equation 15 to obtain an estimate for this static localization error (Table S2). We then subtracted this error from the MSDs to remove the apparent sub-diffusion in the log-log scale plots, this is shown in Figure S10. Notice that for the earliest time lags the dynamics appear to be diffusive, this is the effect of the dynamic localization error on the sub-diffusive trajectories. From the fit, we determined that *α* ≈ 0.75 for the larger particles and *α* ≈ 0.45 for the 20-nm particles.

#### S3. Determining nucleoid occupation time

To calculate the nucleoid occupation time we developed custom code in Matlab. For each cell, we segmented the nucleoid from the DNA label *z*-stack as follows. First, we selected the stack with the best focus of the nucleoid, and then applied an edge detection using the built-in-edge function in Matlab. We used the ‘log’ method for the edge detection function which detects edges by looking for zero-crossings after filtering the image with a Laplacian of Gaussian (LoG) filter [17]. The detected outline is then shrunk by 1 pixel, to account for the blurring from diffraction. The resulting outline is taken to be the shape of the nucleoid in both the *xy* and *xz* cross-sections. We then check the trajectory points, to determine if they fall inside or outside the polygon surrounded by the nucleoid outline. Finally the points are classified as inside the nucleoid if they are inside both the *xy* and *xz* cross-sections. Figure S11 shows an example of the nucleoid segmentation (A) and the points classification (B).

#### S4. Colloidal whole-cell model details

##### Determination of size and abundances of cellular components and macromolecules

We chose the radii of the nucleoid region to be 200 nm to reflect the average values measured in SPT experiments in *E. coli*. We constructed a nucleoid via a random self-assembly process, forming an interconnected colloidal dispersion out of polydisperse spheres of average radius *a*_DNA_ = 7 nm. The self-assembly process can be tuned to produce a network structure that recovers the nucleoid morphology measured in prior experiments [18] (Fig. S12). A volume fraction of *ϕ*_n_ = 0.10 was chosen based on the DNA density at rapid growth conditions prior to entry into stationary phase, and the homogenized nucleoid structure, self assembled by isotropic particles interacting with a hard-sphere Morse potential of depth 10*kT*, was found to more closely reproduce dynamics and localization from the experiments in the present study, perhaps due to changes in nucleoid morphology upon entry into stationary phase. The self-assembled porous network was shaved into a spherical shape and placed in the center of the confined domain.

The radius of the cell was chosen to be 383 nm to match the relative nucleoid to cell volume of 0.39, which was determined from experimental images of the stationary phase cells (see Sec. and). Freely-diffusing macromolecules in the cytoplasm were packed into the remaining volume, until the total occupied volume fraction reached *ϕ* = 0.31. The overall and relative abundances of ribosomes and non-ribosomes prior to entry into stationary phase were taken from reported values for *E. coli* growing at a doubling rate of 3 dbl/hr [19], which was the growth rate of the *E. coli* prior to entry into stationary phase. While there are no explicit measurements for relative ribosome and protein concentrations in stationary phase *E. coli*, we used numerous quantitative reports of degradation of macromolecules in *E. coli* to estimate cellular crowding after entry into stationary phase. The total amounts of ribosomes and other native macromolecules, which we term crowders, were hence determined by adjusting for degradation of 50% of ribosomes [20] and degradation of 20% of proteins [21] upon entry into stationary phase. 40% of ribosomes which are made up of ribosomal proteins were added to the total crowder abundance since ribosomes degrade into proteins and rRNA, whereas the rRNA that make up the ribosomes was not added to the cellular crowding since it was shown to degrade upon entry into the stationary phase [22].

We thus included 19,160 positive and 5,227 negative crowders of radius *a*_crowd_ = 7 nm to reflect the relative abundance and charge of proteome in the *E. coli* cytoplasm [23]. Crowders were added to the model to increase overall effective crowding which comes from not only individual proteins but also tRNA, ternary complexes, polymerase, ribosomal subunits, and other macromolecules in the cell. The size of 7 nm was chosen because it was determined that this size is sufficient to induce size-based demixing when paired with larger macromolecules like ribosomes [24]. We also included 2,931 ribosomes of size *a*_rib_. We represent 20% of ribosomes as part of polysomes – linear chains of six ribosomes [25] – to reflect abundances translating ribosomes in stationary phase *E. coli* [26, 27]. In each simulation, we place 50 GEMs of size *a*_GEM_ = 10 nm, 20 nm, and 25 nm, as in the SPT experiments. This number was chosen as it provides adequate statistics for localization and MSD but is low enough such that we observe no clustering of GEMS in our simulations. All parameters used in whole-cell model construction can be found in Table S5.

##### Macromolecular interactions

A pair of molecules *i* and *j* can interact in the whole-cell model via entropic or electrostatic components, defined by the potential:

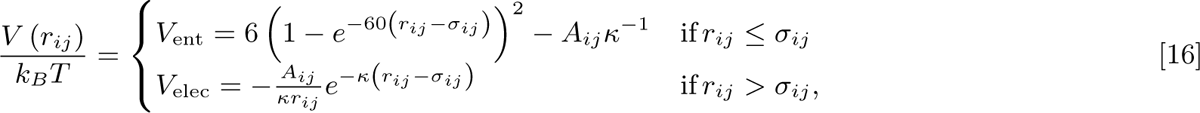

where *r_ij_* is the magnitude of the vector pointing from the center of particle *j* to the center of particle *i*, *k_B_* is the Boltzmann constant, and *T* is the absolute temperature. Entropic exclusion is enforced via a Morse potential, *V*_ent_, which has been previously demonstrated to recover hard-sphere behavior [28]. The hard-sphere contact distance *σ_ij_* is defined as the sum of the particle radii, *a_i_* and *a_j_* (see Table S5).

We modeled electrostatic interactions via a Debye-Hückel potential [29], *V*_elec_, where *κ^−^*^1^ is the Debye length that sets the range of the screened electrostatic interactions. In our model, we chose a value of *κ^−^*^1^ = 2.2 nm, as estimated in the *E. coli* cytoplasm [23]. The strength of the electrostatic attraction or repulsion *A_ij_* is calculated for each pair *i, j*, according to their charges:

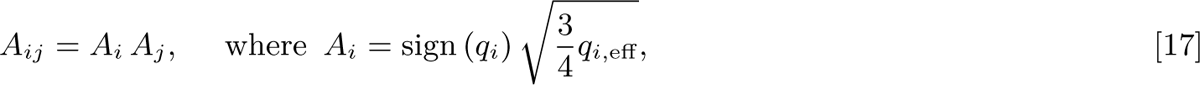

*q_i_* is the nominal charge, and 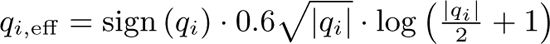 is the effective charge of molecule type *i*. These coarse-grained electrostatic interaction potentials were derived from and parameterized by all-atom simulations; for more detail please see [29]. In our model, pairs thus experience either electrostatic attractions or repulsions depending on their relative charges (Fig. S13). To match experimental conditions, we simulated three types of charged GEMs, where individual GFP monomers have charges of 18*e*, 7*e*, and +15*e*. With 120 monomers per GEM, the total GEM charges are *q*_GEM_ = − 2160*e*, 840*e*, and +1800*e*, respectively. The 20 nm particles have a charge of 420*e* and the 50 nm particles were assumed to have a charge of 1050*e* We assumed each ribosome has a total charge *q*_rib_ = − 3966*e*, as calculated for *M. genitalium* ribosomes in [29]. Each DNA bead has a charge *q*_DNA_ = − 616*e*, corresponding to a surface charge density of 150 mC*/*m^2^ = −1*e/*nm^2^ [30]. We set the charge of positive native crowders to be *q*_crowd,pos_ = +34*e* and negative native crowders to be *q*_crowd,neg_ = − 70*e* based on charge densities of 0.056*e/*nm^2^ and 0.114*e/*nm^2^, respectively. These charge densities reflect the average value for positively and negatively charged proteins in the *E. coli* proteome [23].

Neighboring ribosomes within polysomes – linear chains of six ribosomes each – were connected by a harmonic bond of the form:

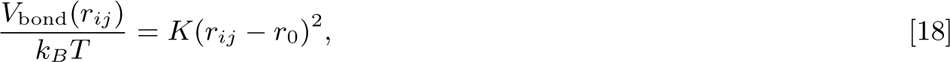

where *K* = 50 sets the bond stiffness and *r*_0_ = 1.08 *σ*_rib_ sets the equilibrium bond distance beyond hard-sphere contact, which prevents frequent steric encounters between neighboring ribosome beads. Polymeric dynamics were enforced for the DNA with a harmonic bond as well, with a bond stiffness of *K* = 200 and *r*_0_ = 1.0 *σ*_DNA_. DNA monomers interact with their bonded neighbors only via the stiff harmonic potential, but interact with other molecules enthalpically and entropically.

Spherical confinement was enforced for all macromolecules *i* with a harmonic force of magnitude *F ^C^*(*r_i_*) that acts inwards in the radial direction:

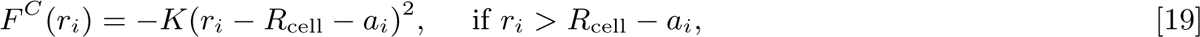

where *K* = 10 and *r_i_* is the distance of the molecule from the center of the cell.

##### Dynamic simulation

The motion of each particle, both the DNA beads and the freely diffusing cytoplasmic molecules, is governed by the Langevin equation, which is a stochastic force balance, given by:

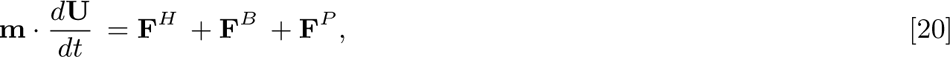

where m is the mass tensor, U is the particle velocity, F*^H^* is the hydrodynamic force, F*^B^* is the stochastic Brownian force, and F*^P^* are forces resulting from particle-particle or particle-boundary interactions.

The hydrodynamic drag on each particle is approximated by the freely-draining limit, meaning the force is calculated using Stokes’ drag law:

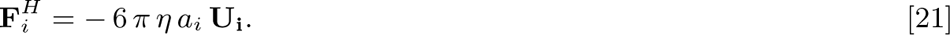

The stochastic Brownian force arises from thermal fluctuations and obeys Gaussian statistics:

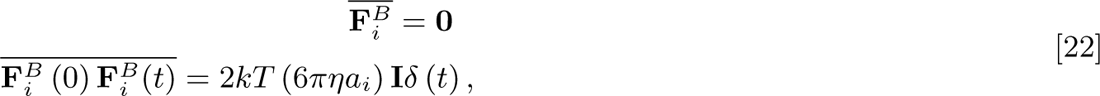

where *k* is the Boltzmann constant, *T* is the absolute temperature, *η* is the solvent viscosity, *a_i_* is the particle radius, I is the identity tensor, and *δ*(*t*) is the Dirac delta function. The overbar considers averages over times that are long compared to the solvent molecule timescales. The interparcle and particle-cavity forces are given by the gradients of the interaction potentials described in described in Section.

Initial configurations for the simulations without GEMs were generated using the software Packmol [31], and the system was equilibrated until a steady-state value was reached for the volume fraction of each species inside versus outside the nucleoid. Following the first equilibration, GEMs were added with the probability of being inside versus outside the nucleoid being determined by experiments. Before data was collected, a second equilibration, this time with GEMs, was performed until GEM localization reached a steady state. The timestep was chosen to be 42 ps to maintain inertia-free dynamics. Simulations were evolved for 3.4 ms, with particle trajectories output every 0.8 *µ*s. Visualizations of the model were created using Visual Molecular Dynamics (VMD, [32]).

#### S5. Colloidal simulation analysis

We calculated the probability distribution function of void segment widths in our self-assembled colloidal gel nucleoid using a radical Voronoi decomposition method developed in our group [33]. We used this method to compare the structure of the model porous network to experimental visualizations of the *E. coli* nucleoid under different conditions (Fig. S12), and compute the fraction of the void distribution accessible to molecules of a given size (Fig. 2D).

To compute the nucleoid occupation likelihood, *t*_nucl_*/t*_tot_, in simulation, we measured the fraction of GEMs located inside of the nucleoid at each time point and averaged this value over 1.2 ms following equilibration. Please see S3 for details regarding how nucleoid occupation was obtained in experiments.

We monitored dynamics of GEMS by tracking particle positions throughout simulation and computing their mean-square displacement (MSD) over time

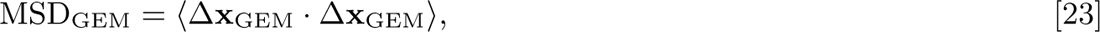

where Δx = x(*t*) − x(0) is the change in particle position from the initial time to the current time *t*. Angle brackets denote an ensemble average over all GEMS. We compare dynamics inside versus outside the nucleoid by separating individual particle trajectories into two different groups: trajectories that take place inside versus outside the nucleoid.

To quantify GEM-macromolecule binding, we calculated the number of molecules of type *i* to which each GEM was bound – its coordination number – and computed the distribution of contact numbers across the GEM ensemble. A GEM, *j*, is considered bound to a molecule, *i*, if their hard sphere surfaces are within 1 nm of contact. We calculated contact number distributions for all other molecules *i*: ribosomes, DNA, and positive and negative crowders (Fig. S14 S14). Using this data, we calculated mean coordination number between a GEM and each molecule type and normalized this by the mean coordination number for a GEM with no net charge (Fig. 3d).

#### S6. Probabilistic simulation of excluded volume nucleoid

Long-time probabilistic simulations of GEM diffusion were performed to explore the roles of confinement and geometry on anomalous diffusion. In these simulations, the shape of *E. coli* was modeled as a cylinder with spherical caps, while the nucleoid was modeled as a smaller cylinder at the center of the cell. Previous methods employed to generate coarse-grained random walks in *E. coli* involve either complete exclusion from the nucleoid [34] or homogeneous diffusion within the entire cell [35]. We expand these approaches to be able to incorporate heterogeneous diffusion (different diffusion coefficients in the nucleoid and in the nucleoid-free region) and non-uniform entry/exit probabilities from the nucleoid, which enable us to reproduce nucleoid dwelling times from the experiments.

Each GEM has a volume normalized tendency to be inside or outside the nucleoid given by 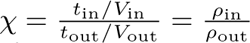, where *ρ*_in *t*_out*_/V_*out *_ρ_*out and *ρ*_out_ are the number densities of GEMs inside and outside the nucleoid given ergodicity of the system. Here, a value of *χ >* 1 indicates preferential localization inside the nucleoid and less than indicates outside. If *χ* = 0, all GEMs are excluded from the nucleoid, and infinitely large values of *χ* indicates confinement within the nucleoid region. Further, we define the ratio of diffusion coefficients inside and outside the nucleoid as *δ* = *D*_in_*/D*_out_. Using these parameters, we employ the following random-walk algorithm that represents heteregeneous dynamics and nucleoid localization times:

1. Initialize *N_i_* GEMs to be inside or outside the nucleoid, with the volume normalized bias toward the nucleoid given by *χ*.
2. For each GEM *i*, generate a displacement in each of the cartesian coordinate directions from a Gaussian distribution of zero mean and variance 2 *D_i_* Δ*t*, where *D_i_* is the diffusion coefficient given by its current location inside or outside the nucleoid and Δ*t* is the time step of the simulation.
3. If the move causes the GEM to go outside of the cell, reject the move and keep the GEM in its original position.
4. If the move would cause the GEM to go from inside the nucleoid to outside, accept the move with a probability given by *p*_out_, which indicates the probability of the GEM escaping the nucleoid region. Similarly, if a GEM goes from outside to inside the nucleoid, accept the probability with *p*_in_. While *p*_in_ and *p*_out_ are parameters that can be varied to represent different physical scenarios, in order to ensure that migration into and out of the nucleoid happen at the same rate (to preserve the initial relative number of GEMs inside and outside the nucleoid), the following relation must hold: 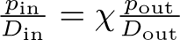
5. Accept all other moves.
6. Repeat for the desired number of time steps.

The values of *p*_in_ and *p*_out_ determine the degree of confinement within the nucleoid or exclusion from the nucleoid. For each GEM, we ran simulations different scenarios to see if experimental MSDs were recovered, as shown in Fig. S16. Free diffusion between regions, with *p*_in_ = 1 if *χ >* 1 and *p*_out_ = 1 if *χ <* 1 alongside the relation 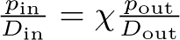 is labeled in Fig. S16 as “porous sphero-cylindrical shell”. Another mode of diffusion that was simulated was tw*^D^*o-^in^region*^D^*c°o^ut^nfinement, where each GEM is confined to inside or outside the nucleoid depending on its initial location (*p*_in_ = *p*_out_ = 0). Finally, we also simulated the scenarios where all GEMs were initially located and confined to the space inside or outside the nucleoid.

We rigorously tested the above algorithm to ensure that it reproduces the desired relative time spent inside the nucleoid. Furthermore, we benchmarked the algorithm for time step sensitivity, beginning with a time step much smaller than a GEMs Brownian time *a*^2^ */D* and found that time steps up to 10 Brownian times produce the same MSDs. Such large timesteps are feasible since we are only coarse-graining the motion of GEMs over very long lag times and GEMs have no interactions beyond confinement. All macromolecular interactions that influence dynamics are coarse grained to be represented by the diffusion coefficients.

The size of the nucleoid and cell were first determined by experimental images and then adjusted slightly in order to match the plateaus in MSD observed in experiments. Discrepancies between nucleoid and cell sizes determined from experimental images and the random-walk simulations were less than 10%. Diffusion coefficients were varied in order to match experimental MSD data. The ratio of diffusion coefficients inside (*D*_in_) to outside (*D*_out_) the nucleoid were estimated from the colloidal whole-cell simulations by the ratio of MSDs inside and outside the nucleoid at a lag time where particles diffuse an equivalent to one timestep in the probabilitic long-time simulations. For all of these simulations, a bulk diffusion coefficient (*D*_bulk_) was determined from the MSD of all trajectories inside and outside. We also ran simulations with the bulk diffusion coefficient, and found that two-mode heterogeneous diffusion did not produce any discernible characteristics in MSD at this level of coarse-graining, meaning a random walk throughout the cell with *D*_bulk_ produced equivalent MSDs to having separate *D*_in_ and *D*_out_. However, localization statistics that arise from the base penetrability (*p*_in_ and *p*_out_) and ratio of penetration into and out of the nucleoid (*χ*) do impact both the times at which anomolous diffusion emerges as well as the confinement plateaus in MSD.

---

**Fig. S1.**
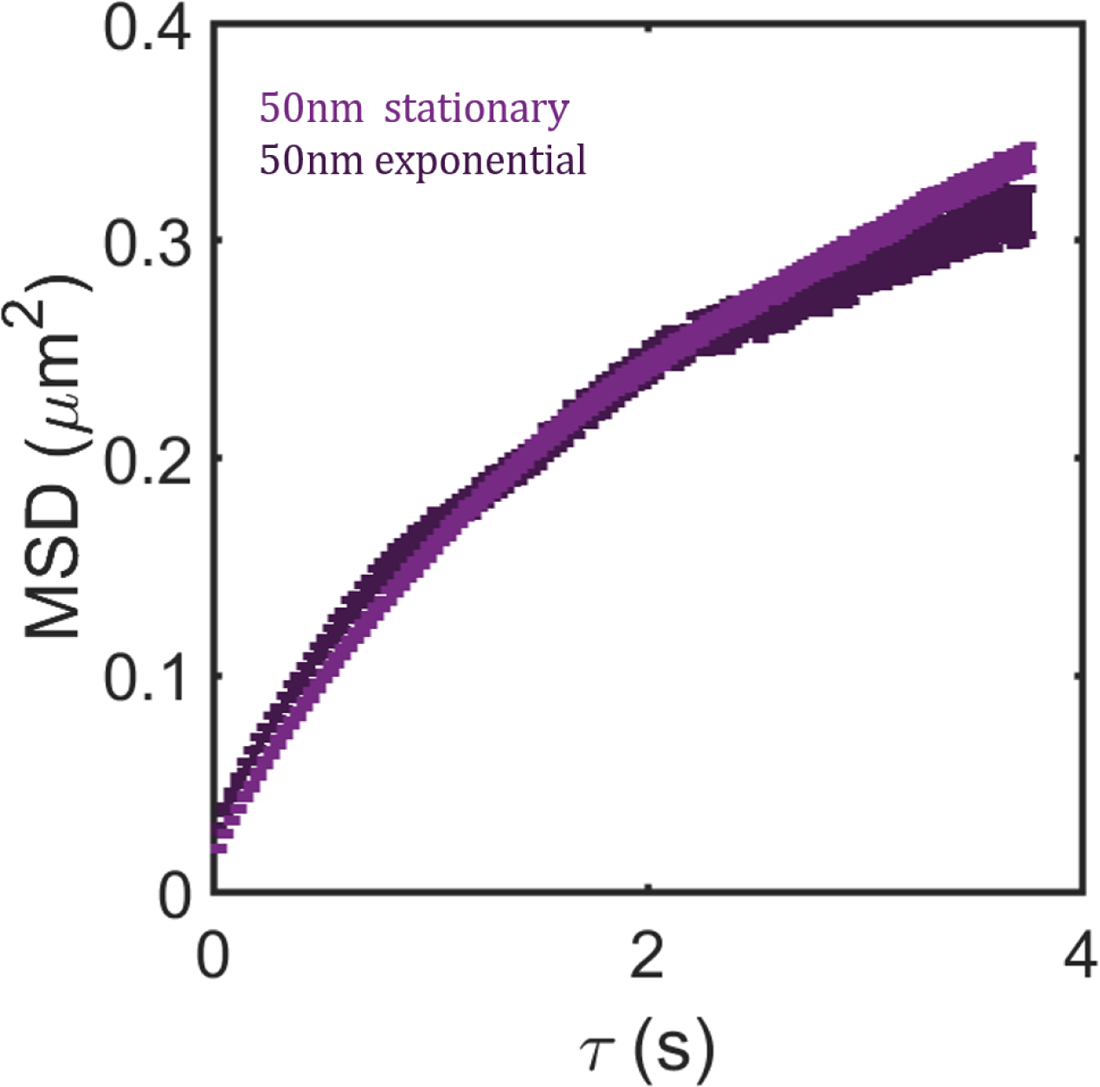
Mean square displacements for 50 nm bGEMs diffusing in exponential and stationary phase cells

**Fig. S2.**
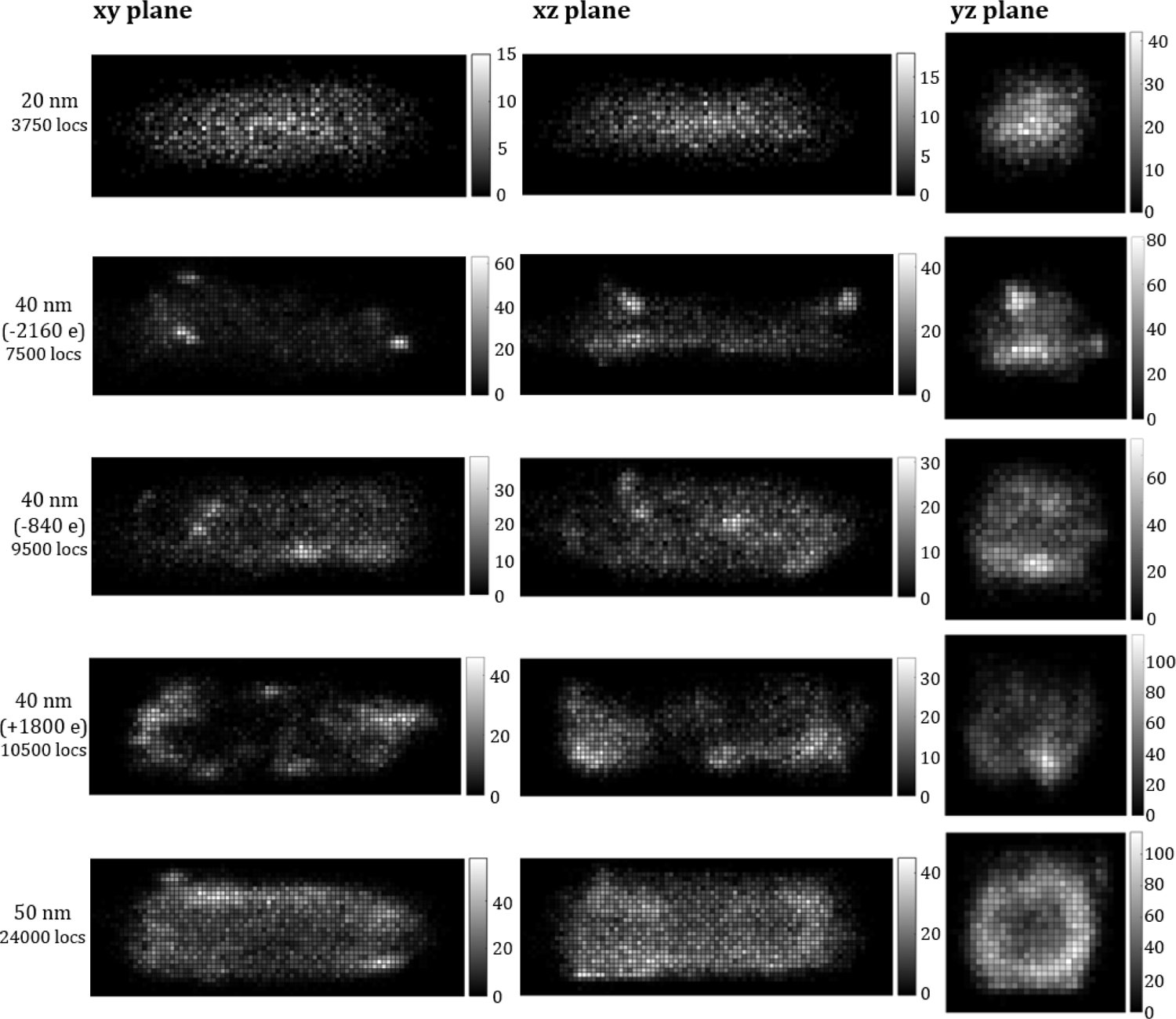
Histograms of particle localization for all bGEMs sizes and charges. Data is shown for all three projection planes *xy*,*yz*,*xz*. The total number of data points for each figure is shown in the text on the left. The data was binned such that each pixel in the images is close to 30nm.

**Fig. S3.**
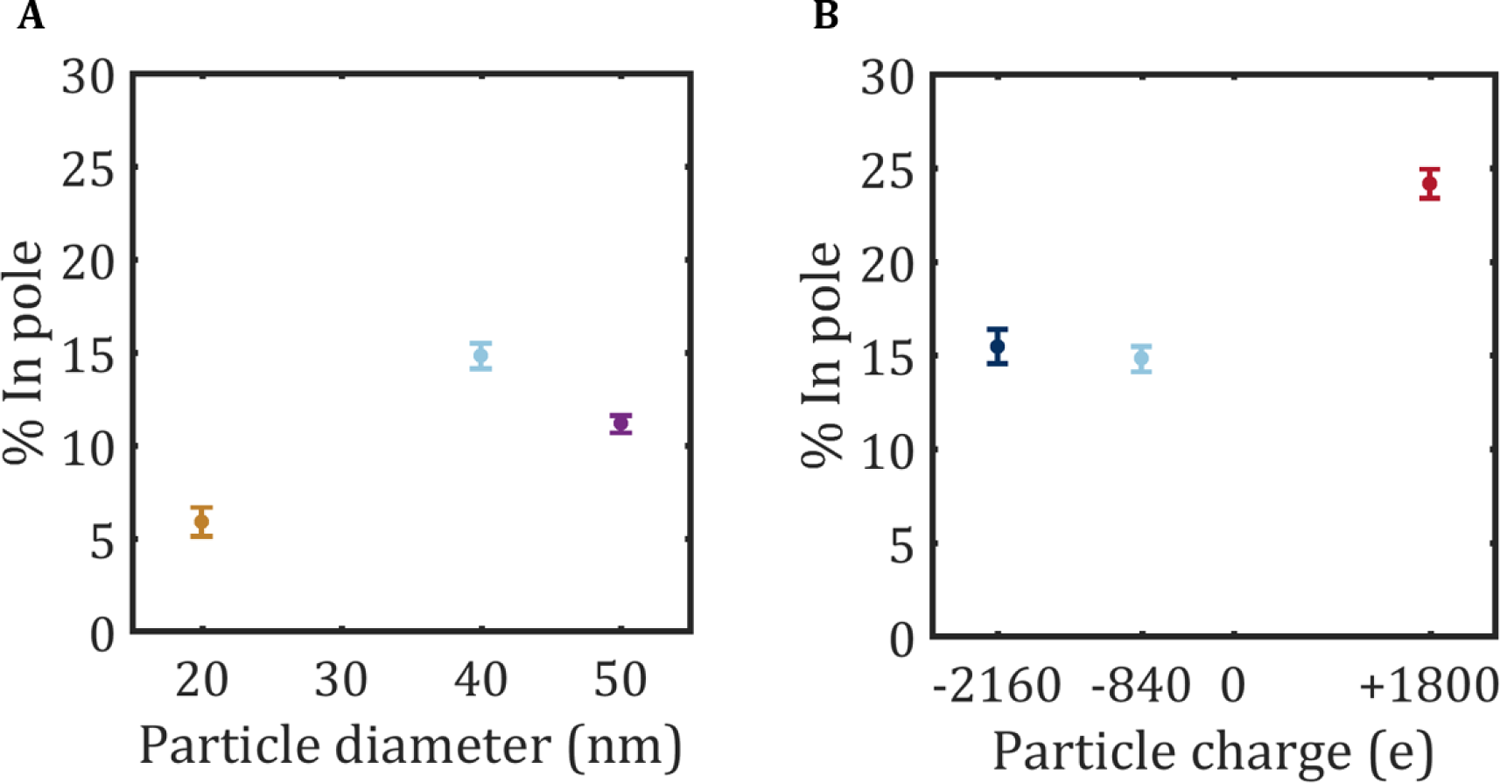
Pole occupation for bGEMs. (A) Percentage of localizations found at the cellular poles for the different size variants. (B) Percentage of localizations found at the cellular poles for the different charge variants.

**Fig. S4.**
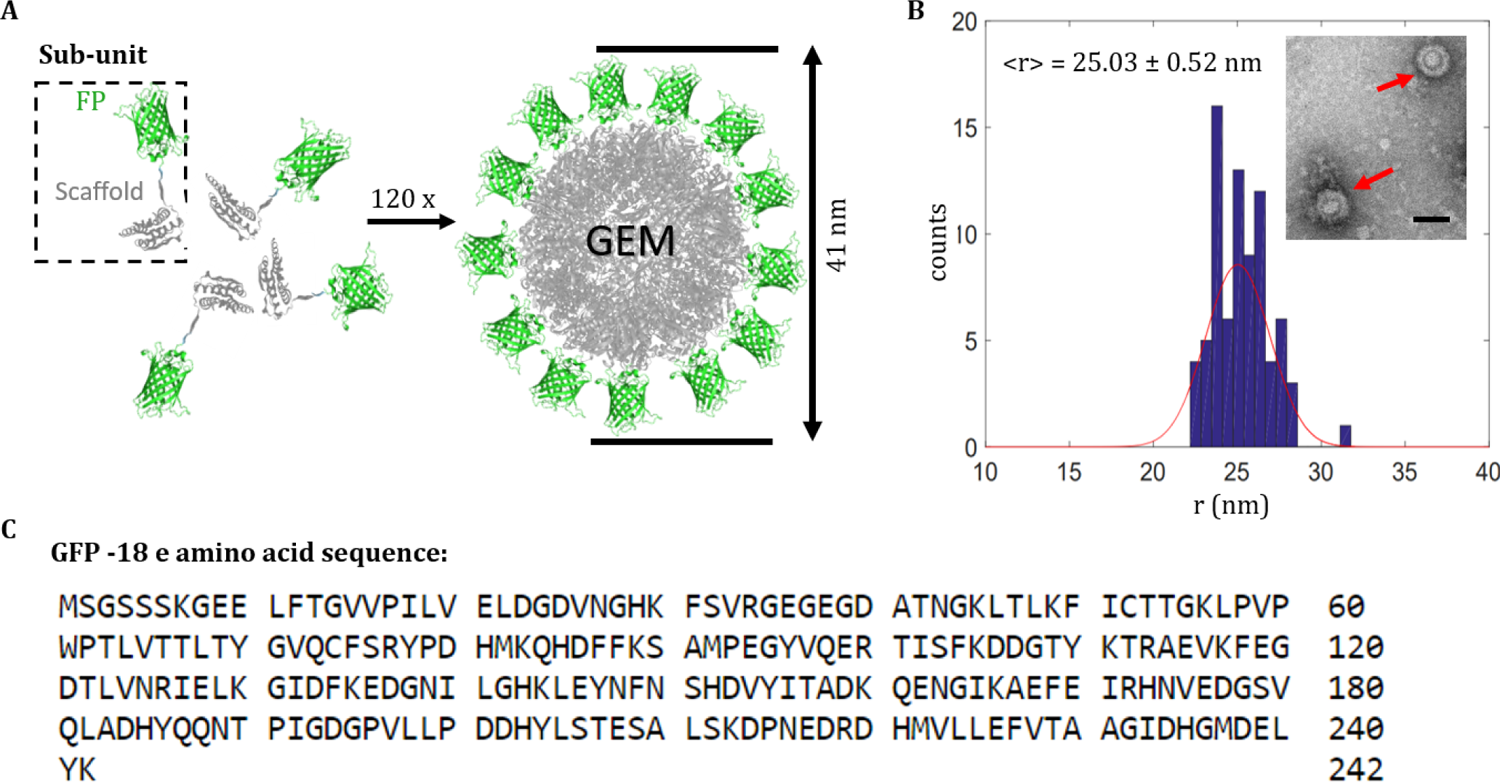
Bacterial Genetically Encoded Multimeric Nanoparticles. (A) bGEMs self-assemble into icosahedral particles from sub-units expressed by a plasmid in the bacterial cell. Each sub-unit consists of a scaffold protein linked to a fluorescent protein (FP). Protein visualizations from[36]. (B) Negative stain transmission Electron microscopy results for new 50 nm GEMs. Inset: TEM image showing 2 50 nm GEMs (red arrows), scale bar: 50 nm. (C) Amino acid sequence for new −18 GFP charge variant.

**Fig. S5.**
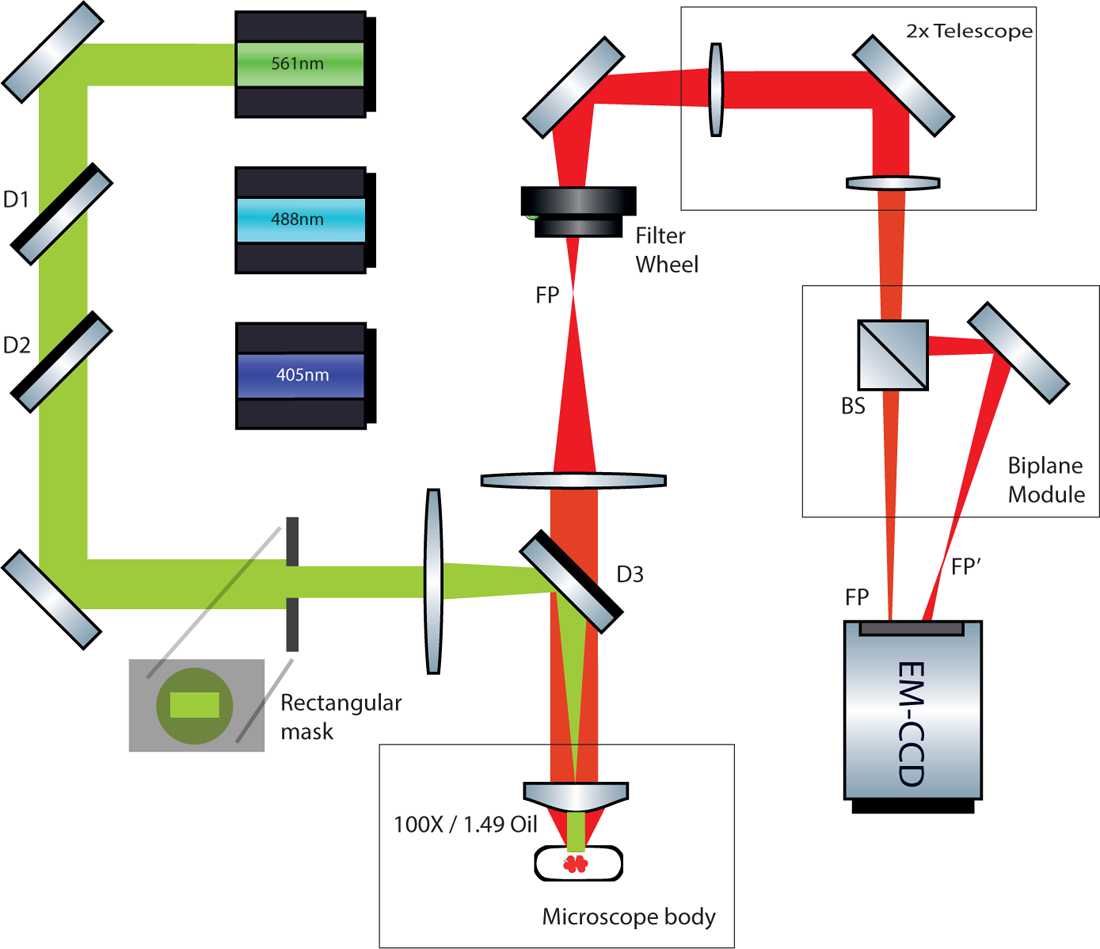
Simplified diagram of the optical setup for the biplane microscope. The biplane module consists of a beam splitter and mirror. This simple addition enables the user to image two separate focal planes simultaneously. On the illumination path, a rectangular mask is used to block part of the beam to create the right shape for illumination in the biplane setup.

**Fig. S6.**
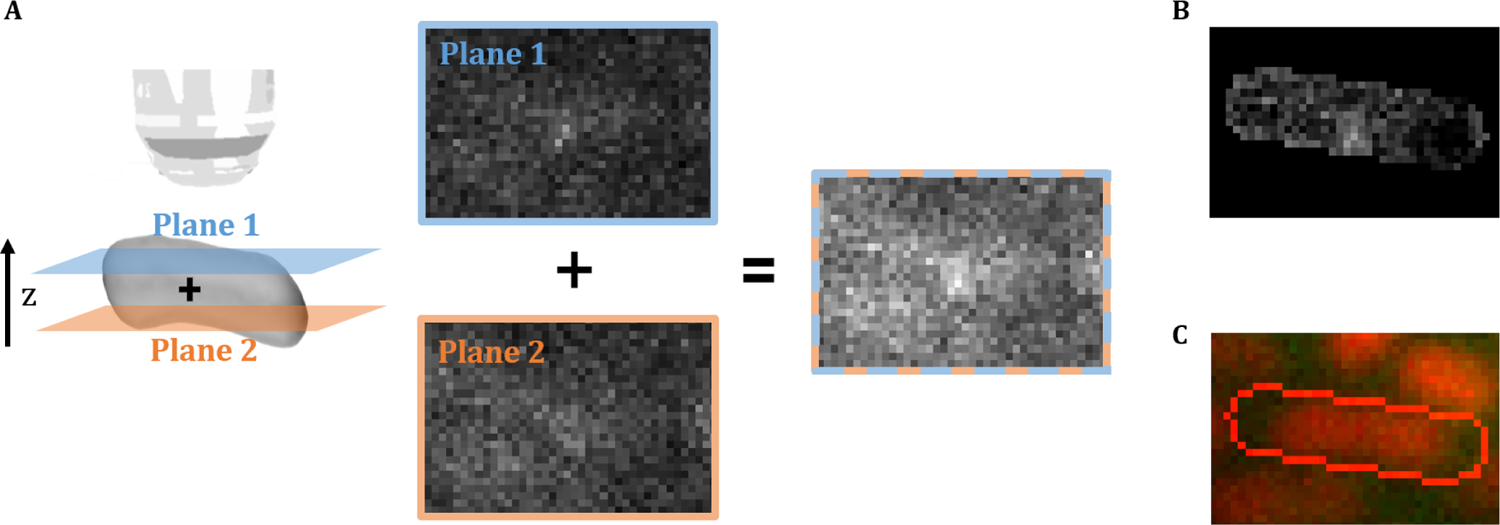
Image analysis pipeline. (A) To create a composite image for tracking the two focal planes are added together. (B)Segmented cell with background subtraction (C)Cell outline overlaid over nucleoid image and GEM channel.

**Fig. S7.**
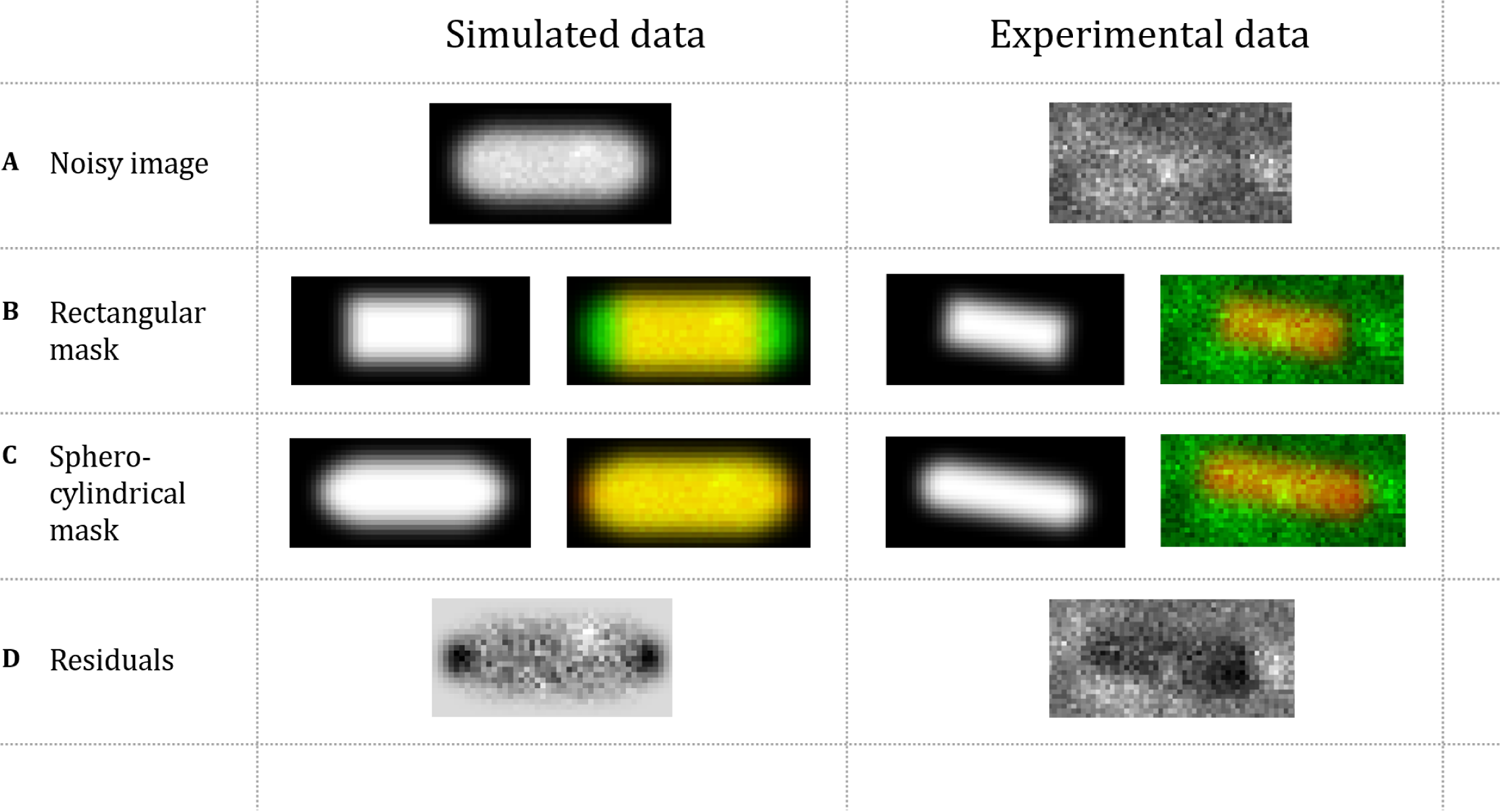
Fitting pipeline exemplified on simulated data and experimental data. Noisy images (A) are first fitted with a rectangular mask (B), then an empirical spherocylindrical mask (C) is fitted to the noisy image. Spot finding is done in the residuals image (D).

**Fig. S8.**
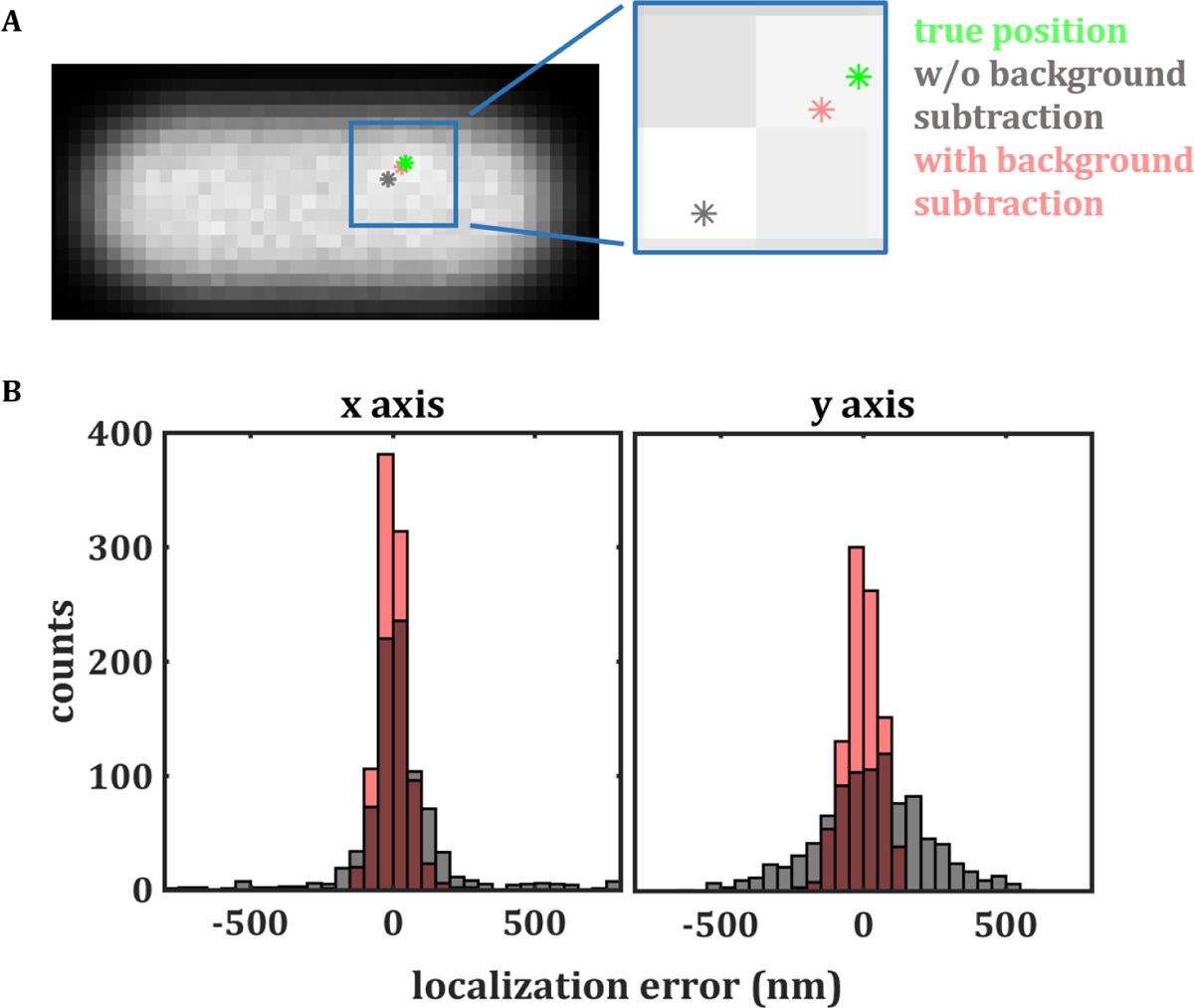
Localization error and background subtraction. (a) Simulated image used for testing the *xy* peak finding algorithm and background subtraction. The background subtracted peak finding (salmon) is closer to the true particle position (green) than the peak finding result without background subtraction (gray). (b) The localization error histograms for the *x* and *y* axis peak finding show a marked improvement when background subtraction is implemented before peak detection.

**Fig. S9.**
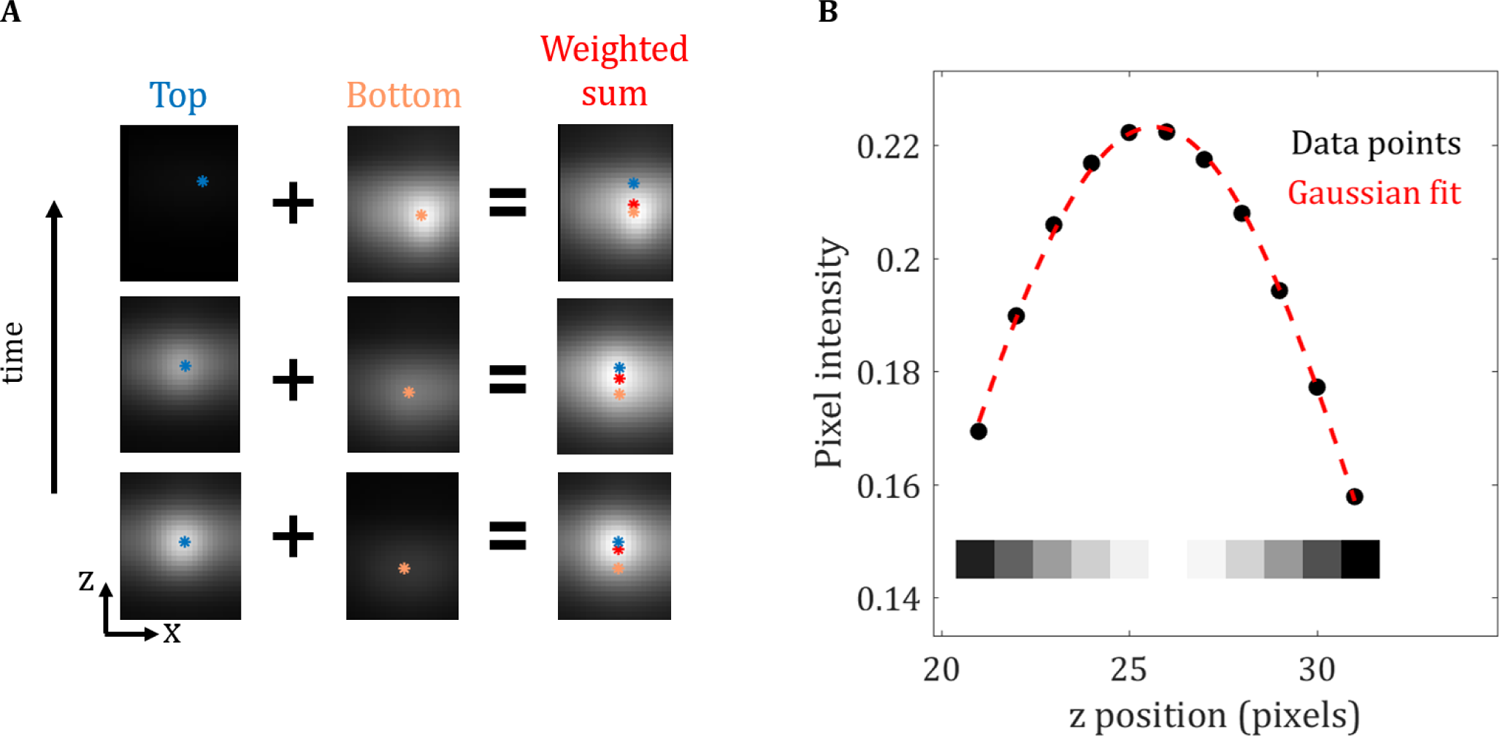
Graphical representation of the convolution tracking algorithm (A) *z*-positions are determined for each plane, a weighted sum of these positions based on the relative focus of the particle is used to determine the true location. (B) Gaussian fit of the intensity profile for a *z*-sliver (10×1) of data surrounding the intensity peak (sliver is taken at the *xy* location of the candidate particle).

**Fig. S10.**
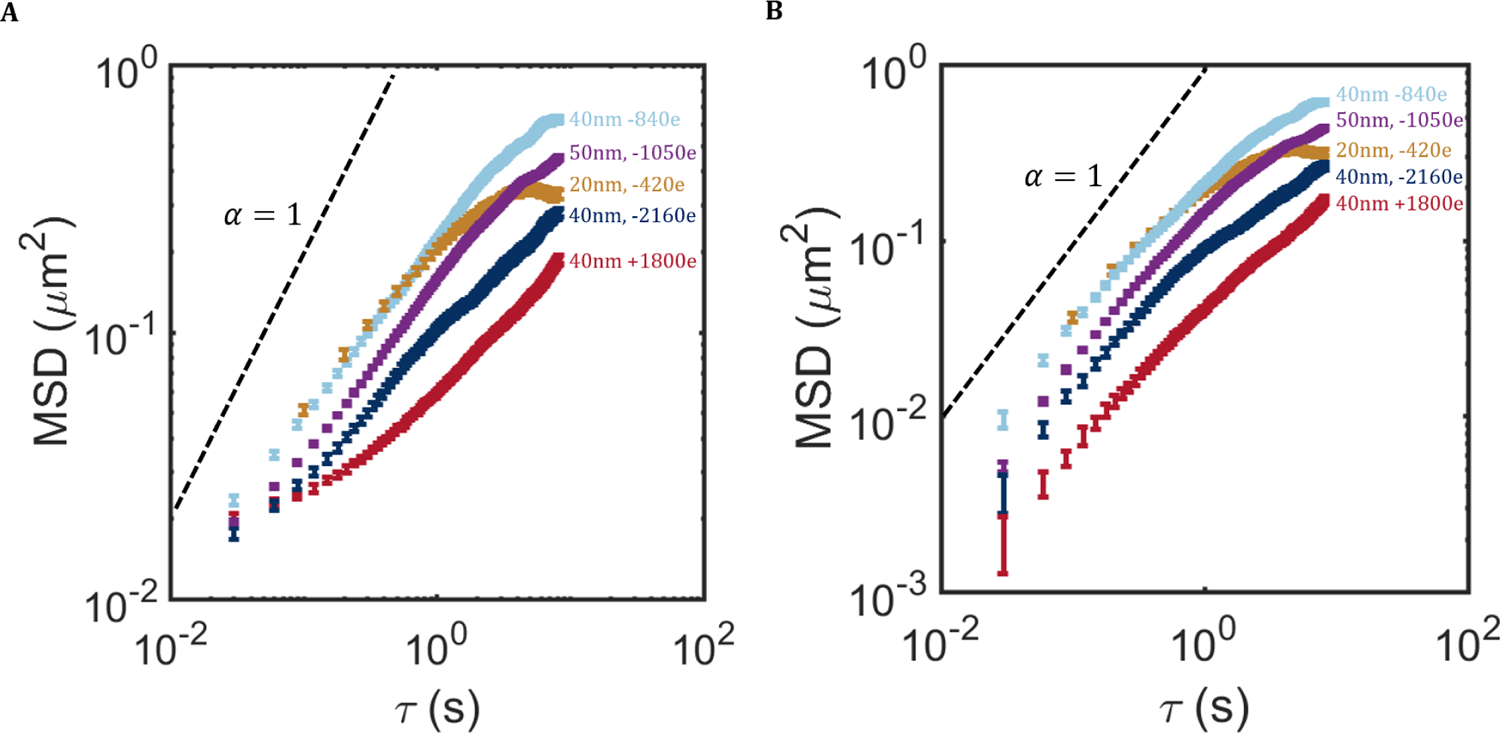
Artifacts in the MSD from static and dynamic localization errors. (A) Shows the uncorrected MSDs, the trajectories appear sub-diffusive at short time lags, this is a result of the static localization error. It appears as though there are multiple diffusion regimes, specially for the 40nm +1800e GEMs. (B) When the static localization error is subtracted from the trajectories, these spurious diffusion regimes disappear. For the first few time lags, the particle’s dynamics appear to be diffusive, this is an artifact of the dynamic localization error.

**Fig. S11.**
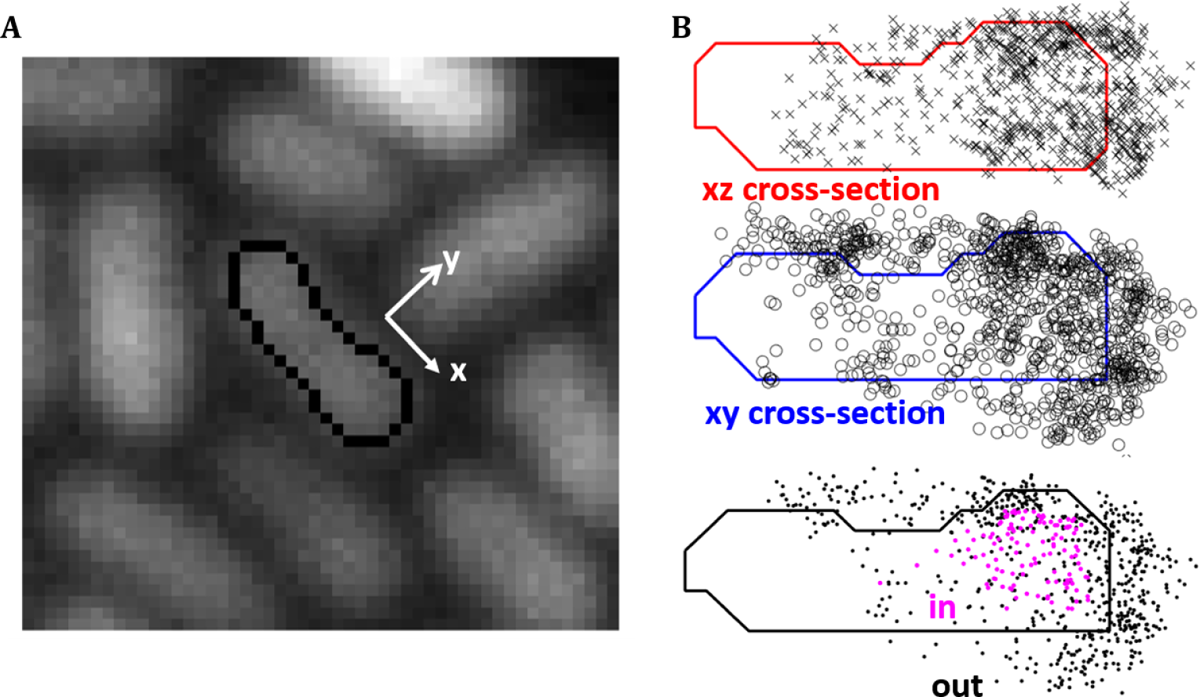
Nucleoid segmentation and trajectory points in nucleoid classification (A) Nucleoid image with a single nucleoid highlighted in black (B) Only trajectory points located inside both nucleoid cross-sections (xy and xz) are classified as inside the nucleoid region. Cross-sections are found from the segmentation shown in A.

**Fig. S12.**
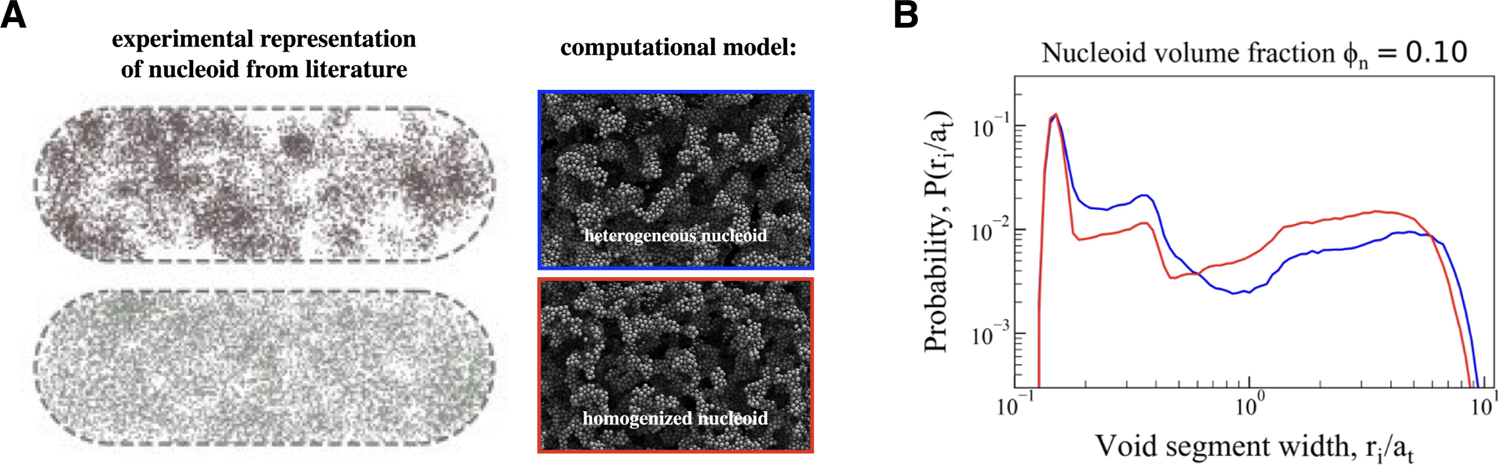
Self-assembled network of colloidal particles represent the *E. coli* nucleoid under different solvent conditions. (A) (left) Visualization of *E. coli* nucleoid morphologies under two different solvent conditions [18]. (right) We construct model nucleoids in simulation that recapitulate both the density of DNA and the nucleoid porosity reported in experiments. (B) Probability distribution function of void radii *ri* [33] for the two model nucleoids reveals that both structures have a heterogeneous distribution of pore sizes, but the bottom (outlined in red) structure is relatively more homogenized. The x-axis is normalized by the characteristic size of a DNA bead, which in this case is *ai* = 7 nm. The red curve represents the pore-size distribution of the nucleoid used throughout the simulations in this work.

**Fig. S13.**
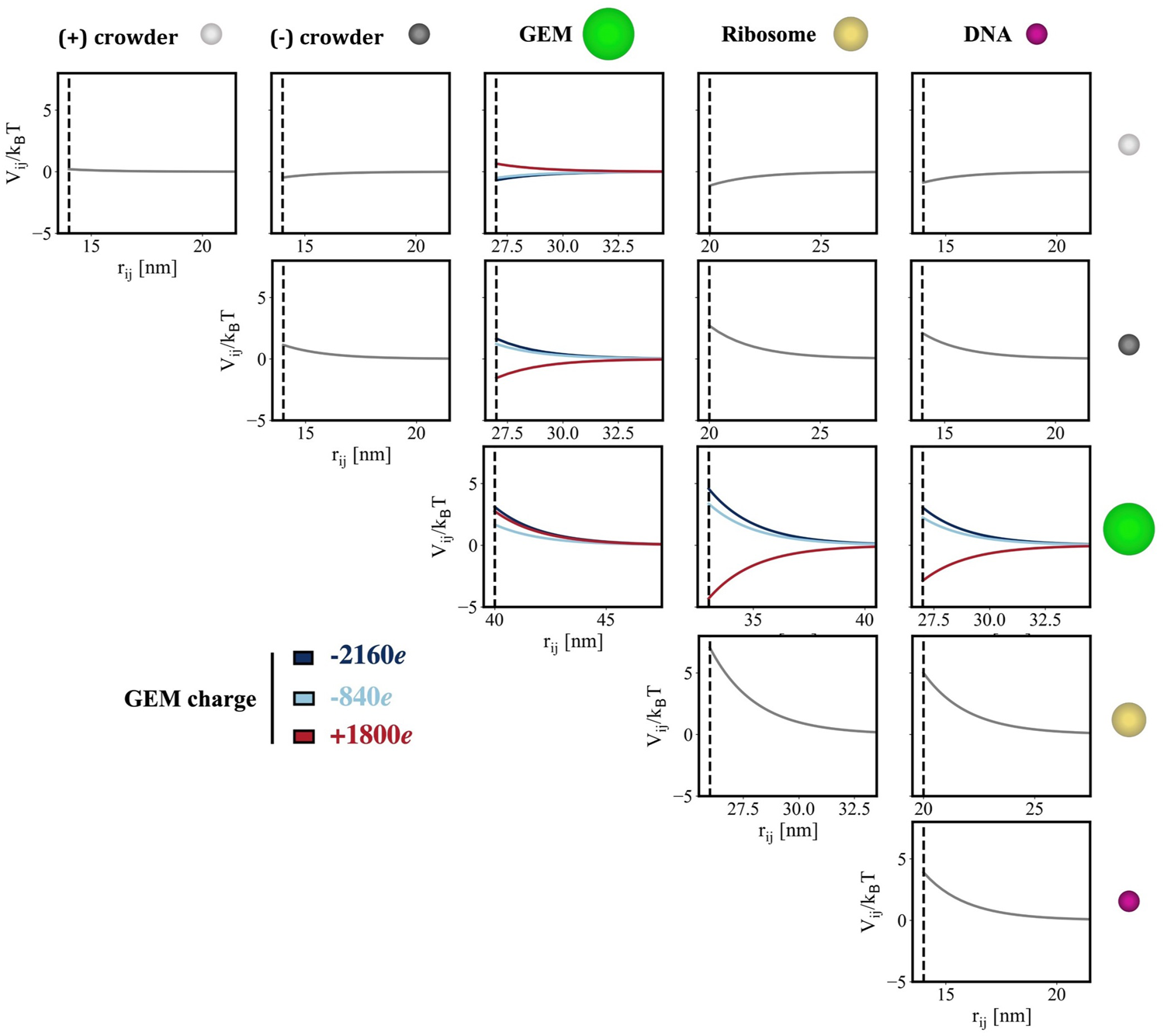
Macromolecules experience electrostatic attractions or repulsions depending on their relative charges. Interaction potentials *V_ij_ /k*_B_*T* between all unique types of macromolecule pairs, *i*,*j* (labels above, and representations to right). Particle pairs that are both negative or both positive (e.g., pairs of the same type, along the diagonal) experience repulsions, where *V_ij_ /k*_B_*T >* 0. The charge of GEMs (40 nm, shown here) can be tuned by changing the charge of individual GFP monomers (legend to left; colors), which can induce moderately strong attractions between positively charged GEMs and ribosomes, DNA, and negative crowders (*V_ij_ /k*_B_*T <* 0). When pairs reach their defined contact distance (vertical dashed lines), they experience a steeply repulsive Morse potential that enforces entropic exclusion.

**Fig. S14.**
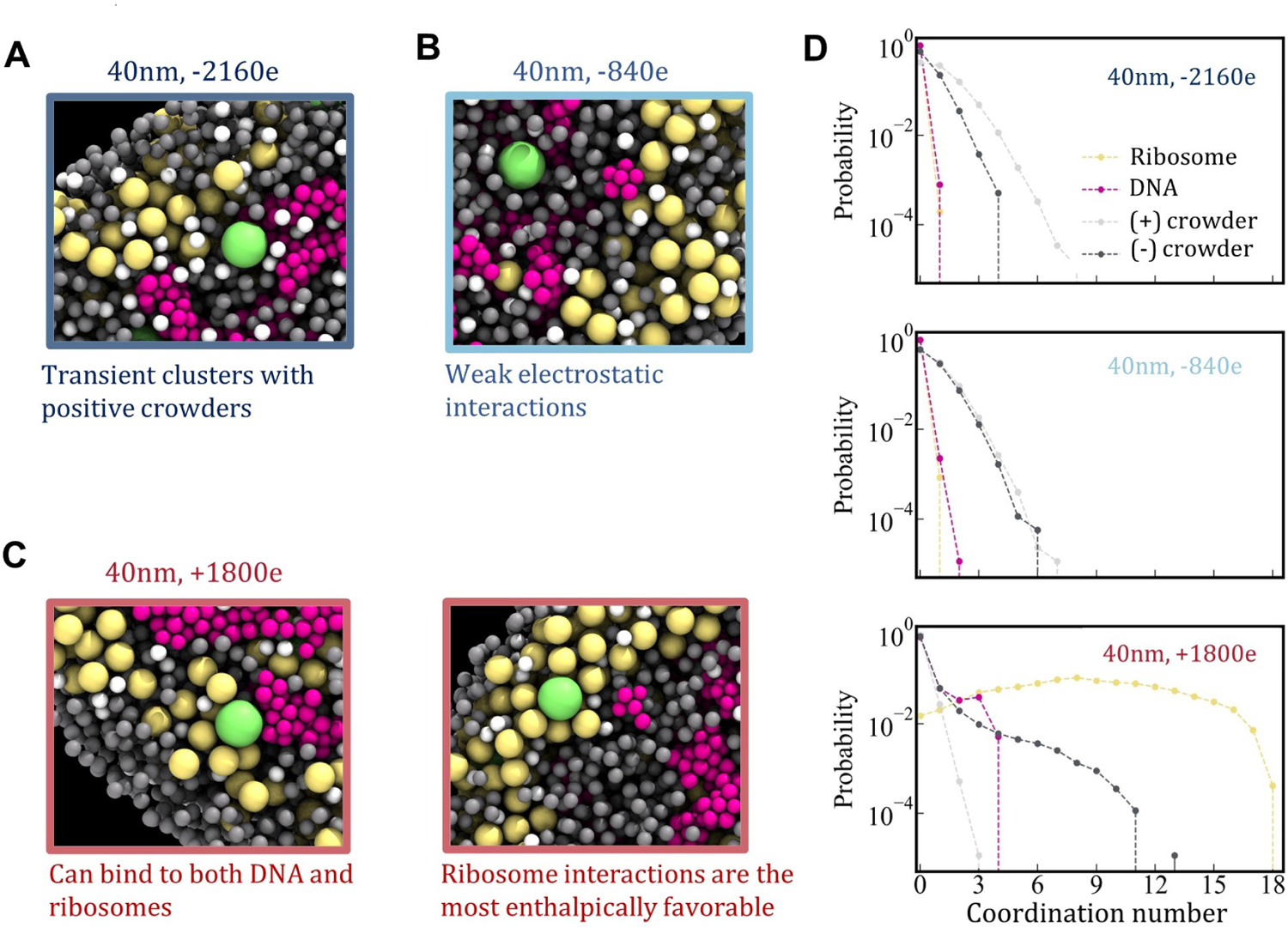
Simulations reveal varying local macromolecular environment as a function of surface charge. Snapshots are from simulations of (A) 40 nm, −2160*e*, (B) 40 nm, −840*e*, and (C) 40 nm +1800*e* GEMs. (D) Probability distribution function of coordination numbers between GEMs of different charges and ribosomes, DNA, and positive and negatively charged crowders quantifies changes in local interactions induced by GEM charge.

**Fig. S15.**
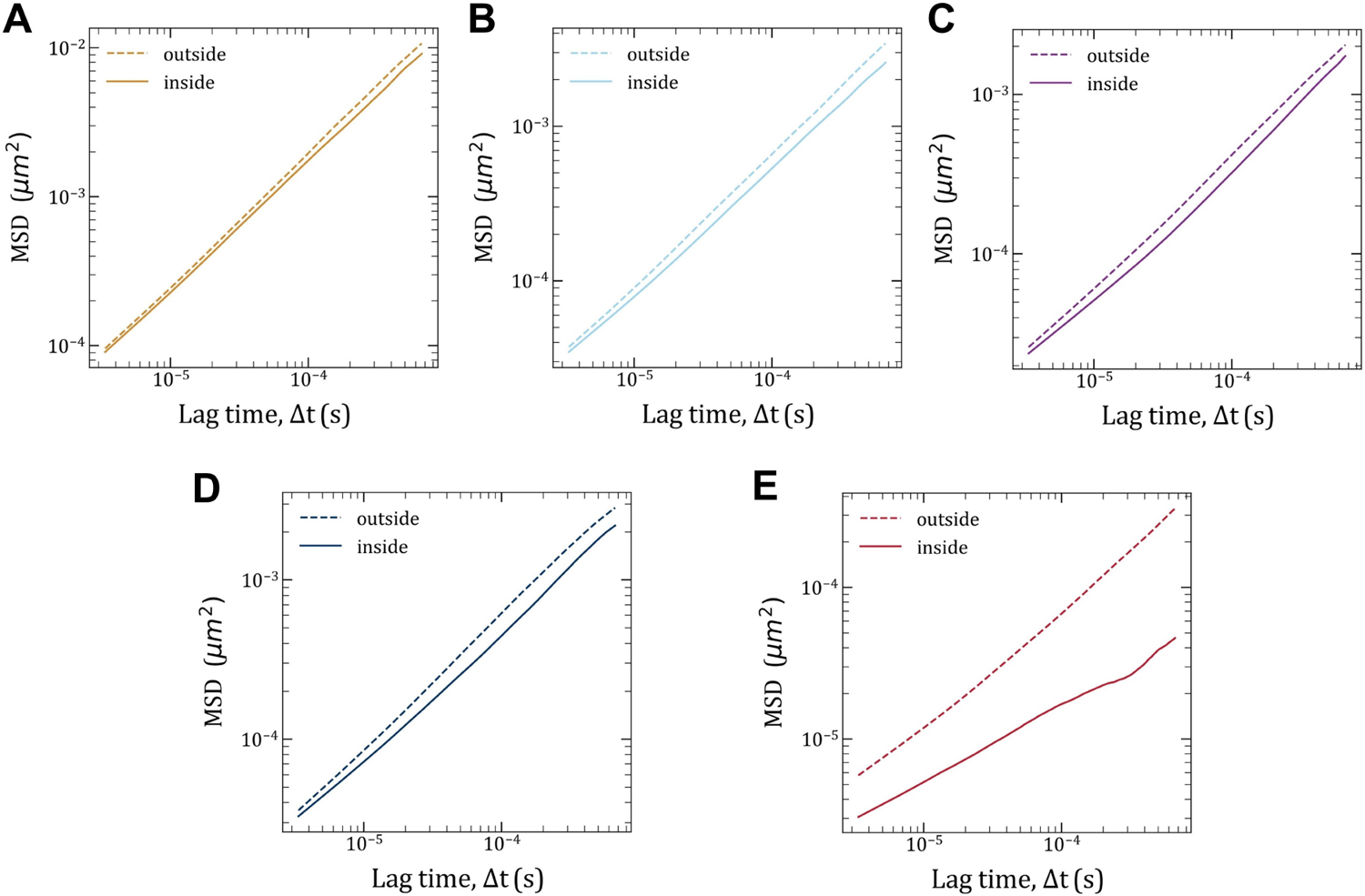
MSD of GEMs inside versus outside the nucleoid obtained from colloidal whole-cell simulations. Data are from simulations of (A) 20 nm, −420*e*, (B) 40 nm, −840*e*, (C) 50 nm, −1050*e*, (D) 40 nm, −2160*e*, and (E) 40 nm +1800*e* GEMs.

**Fig. S16.**
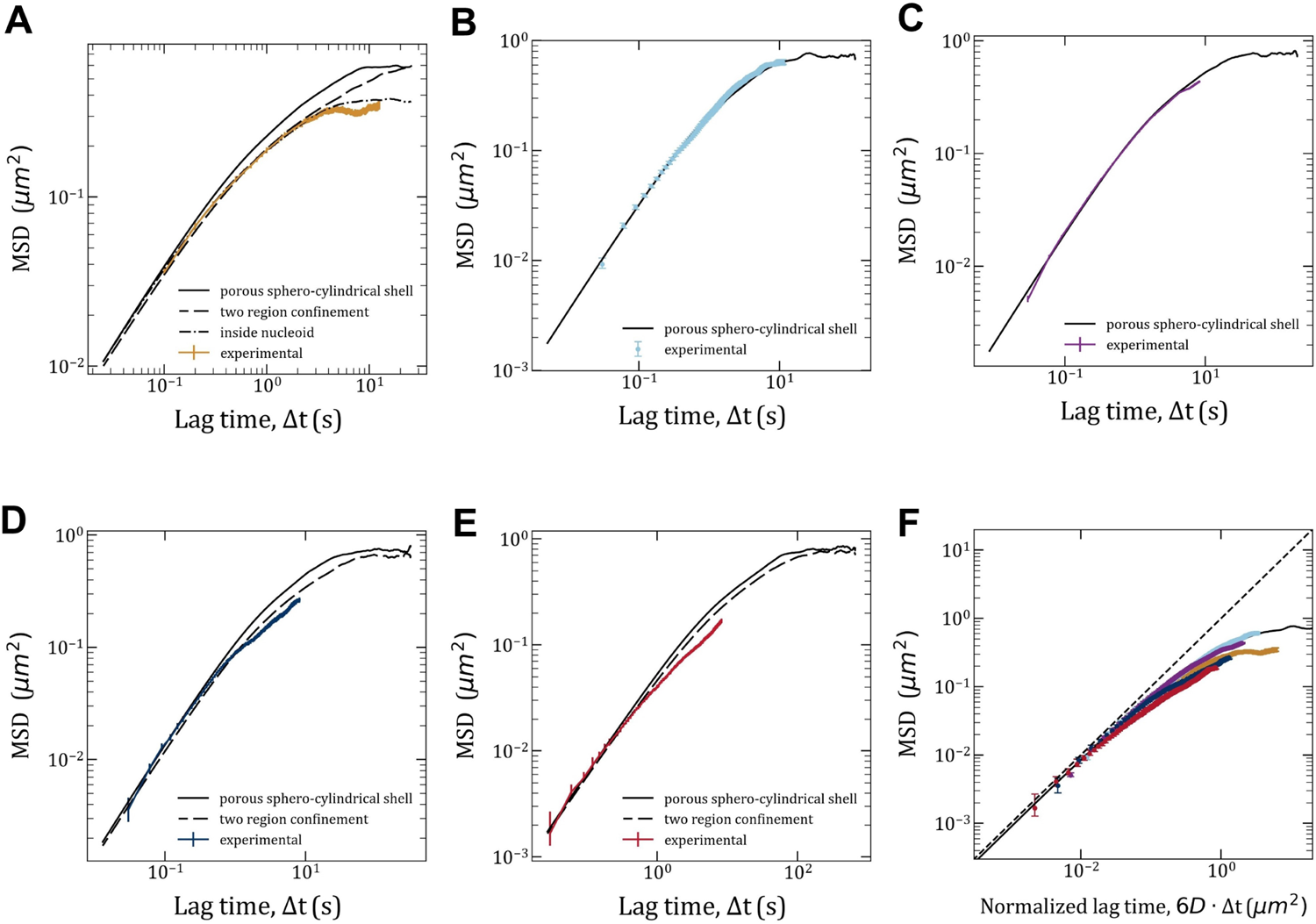
Probabilistic simulation of excluded volume nucleoid reveal the role of geomoetry and confinement on anomolous diffusion in *E. coli*. **(A-E)** MSD over time for (A) 20 nm, −420*e*, (B) 40 nm, −840*e*, (C) 50 nm, −1050*e*, (D) 40 nm, −2160*e*, and (E) 40 nm +1800*e* GEMs. Colored symbols are experimental data. Black lines represent diffusion through a porous sphero-cylindrical shell (solid lines), two-region confinement (dashed lines), or complete localization and confinement within one region of the cell (dotted dashed lines), as described in Sec. Diffusion coefficients and nucleoid occupation times for each simulation are in Table S7. **(F)** MSD versus normalized lag time for all experiments shows that all trajectories display similar dynamic behaviors, with differences in the timescales of confinement plateaus. Solid black line is from simulation in (B) and dotted line is free, unconfined diffusion.

**Table S1.**
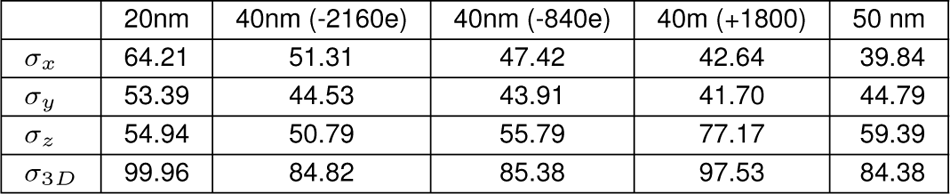
Localization error for each particle size and charge. The errors are obtained from fitting the MSD curves with equation 15. The 1D localization uncertainity is ≈ 50*nm* for all particles.

**Table S2.** Diffusion coefficients and exponents from experiments and simulations. Experimental values are obtained as described in S2. To enable comparison between effective diffusion coefficients a fixed *α* =0.75 was fitted to obtain the diffusion coefficients of particles of diameter 40 nm and 50 nm.

| bGEM | $\alpha_{exp}$ (best fit) | $Deff_{sim}$ | $\alpha_{sim}$ | $Deff_{sim}$ |
| --- | --- | --- | --- | --- |
| 20 nm | 0.43 | $0.040 \mu m^2/s^{0.43}$ | 0.91 | $0.94 \mu m^2/s^{0.86}$ |
| 40 nm (-2160) | 0.68 | $0.015 \mu m^2/s^{0.75}$ | 0.84 | $0.25 \mu m^2/s^{0.86}$ |
| 40 nm (-840) | 0.74 | $0.036 \mu m^2/s^{0.75}$ | 0.88 | $0.29 \mu m^2/s^{0.86}$ |
| 40 nm (+1800) | 0.74 | $0.007 \mu m^2/s^{0.75}$ | 0.86 | $0.024 \mu m^2/s^{0.86}$ |
| 50 nm | 0.78 | $0.024 \mu m^2/s^{0.75}$ | 0.84 | $0.18 \mu m^2/s^{0.86}$ |

**Table S3.**
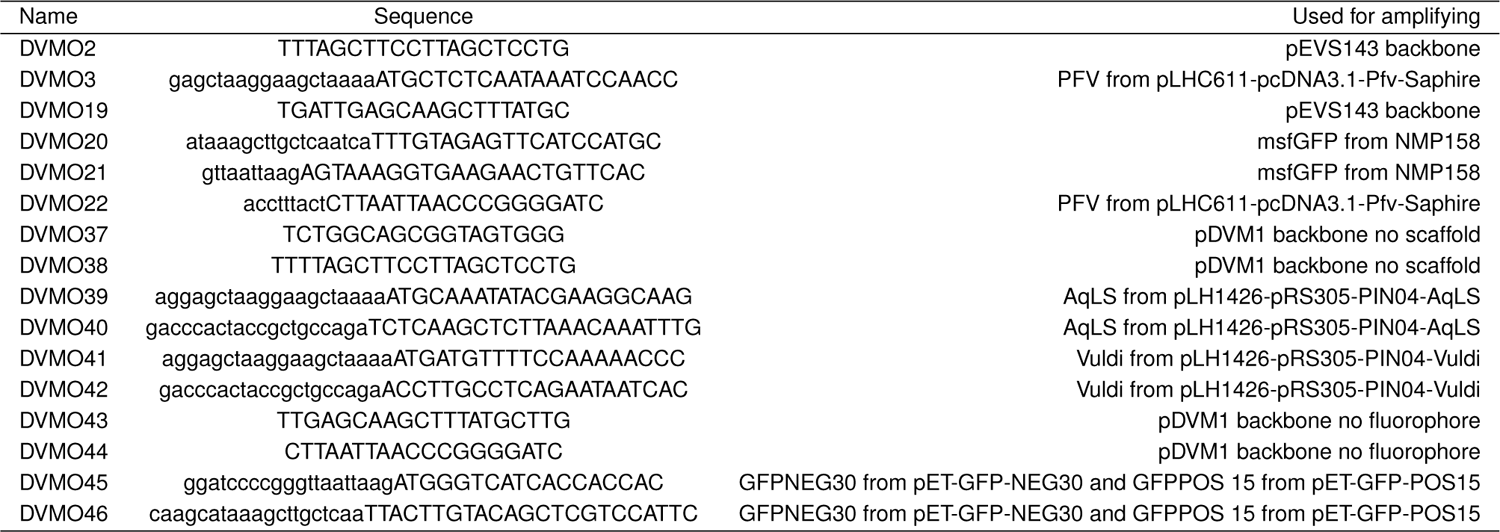
Primers used in this study.

**Table S4.** Strains used in this study.

| Strain | Description | Source |
| --- | --- | --- |
| ZG1352 | WT *E.coli* MG1655 | [37] |
| S17 | Ec used for cloning and plasmid conjugal transfer | [38] |
| DVM085 | MG1655+pEVS143::Ptet-*AqLS-msfGFP*+pSACT101; Kan<sup>r</sup>, Amp<sup>r</sup> | This study |
| DVM076 | MG1655+pEVS143::Ptet-*PFV-msfGFP*+pSACT101; Kan<sup>r</sup>, Amp<sup>r</sup> | This study |
| DVM079 | MG1655+pEVS143::Ptet-*PFV-GFPPOS15*+pSACT101; Kan<sup>r</sup>, Amp<sup>r</sup> | This study |
| DVM083 | MG1655+pEVS143::Ptet-*PFV-GFPNEG18*+pSACT101; Kan<sup>r</sup>, Amp<sup>r</sup> | This study |
| DVM090 | MG1655+pEVS143::Ptet-*Vuldi-msfGFP*+pSACT101; Kan<sup>r</sup>, Amp<sup>r</sup> | This study |

**Table S5.** Simulation parameters for construction of a whole-cell *E. coli* colloidal model.

|  |  |  |
| --- | --- | --- |
| Radius of nucleoid region | $R_n$ | 280 nm |
| Radius of cell | $R_{\text{cell}}$ | 383 nm |
| Volume fraction of DNA within nucleoid | $\phi_n$ | 0.1 |
| Total volume fraction of nucleoid + cytoplasm | $\phi_{\text{tot}}$ | 0.31 |
| Radius of DNA | $\langle a_{\text{DNA}} \rangle$ | 7 nm, $\sigma^2 = 0.5$ |
| Charge of DNA | $q_{\text{DNA}}$ | $-616e$ |
| Number of DNA beads | $N_{\text{DNA}}$ | 6310 |
| Radius of ribosomes | $a_{\text{rib}}$ | 13 nm |
| Charge of ribosomes | $q_{\text{rib}}$ | $-3966e$ |
| Number of ribosomes | $N_{\text{rib}}$ | 2931 |
| Radius of GEMs | $a_{\text{GEM}}$ | 10, 20, 25 nm |
| Charge of GEMs (for 40 nm size) | $q_{\text{GEM}}$ | $-2160e, -840e, +1800e$ |
| Number of GEMs | $N_{\text{GEM}}$ | 50 |
| Radius of positive and negative crowders | $a_{\text{crowd}}$ | 7 nm |
| Charge of positive crowders | $q_{\text{crowd,pos}}$ | $+34e$ |
| Number of positive crowders | $N_{\text{crowd,pos}}$ | 19160 |
| Charge of negative crowders | $q_{\text{crowd,neg}}$ | $-70e$ |
| Number of negative crowders | $N_{\text{crowd,neg}}$ | 5227 |

**Table S6.** Simulation parameters for probabilistic simulations of excluded volume nucleoid.

|  |  |  |
| --- | --- | --- |
| Cylindrical diameter of cell (not including caps) | $l_{\text{cell}}$ | 1400 nm |
| Cylindrical radius of cell | $r_{\text{cell}}$ | 400 nm |
| Radius of spherical caps | $r_{\text{cap}}$ | 400 nm |
| Cylindrical diameter of nucleoid | $l_n$ | 1200 nm |
| Cylindrical radius of nucleoid | $r_n$ | 300 nm |
| Radius of GEMs | $a_{\text{GEM}}$ | 10, 20, 25 nm |

**Table S7.** Parameter and outputs of long-time probabilistic simulations of an excluded volume nucleoid. The ratio *t*_in_*/t*_out_ was obtained from nucleoid occupation times in Figs. 2c and 3c. Diffusion coefficients were tuned to match experimental MSDs (Fig. 4a). Effective viscosity was calculated from the Stokes-Einstein equation with *D*_bulk_ as the diffusion coefficient. For reference, *η*_eff,GFP_ ≈ 9.7 cP in the *E. coli* cytoplasm [39] and *η*_water @ 23 °C_ ≈ 0.93 cP.

| **bGEM Type** | $t_{\text{in}}/t_{\text{out}}$ | $D_{\text{bulk}} (\mu\text{m}^2/\text{s})$ | $D_{\text{in}} (\mu\text{m}^2/\text{s})$ | $D_{\text{out}} (\mu\text{m}^2/\text{s})$ | $\eta_{\text{eff}} (\text{cP})$ |
| --- | --- | --- | --- | --- | --- |
| 20 nm | 1.0 | 0.0791 | 0.0791 | NA | 137 |
| 40 nm, -840e | 0.39 | 0.0674 | 0.0586 | 0.0733 | 80.5 |
| 50 nm | 0.32 | 0.0397 | 0.0324 | 0.0432 | 109 |
| 40 nm, -2160e | 0.41 | 0.0259 | 0.0208 | 0.0296 | 209 |
| 40 nm, 1800e | 0.25 | 0.0122 | 0.00446 | 0.0149 | 445 |

## Notes

### Competing Interest Statement

The authors have declared no competing interest.

### Summary of Updates

Fixing figure 1 caption, original caption was clipped.

